# GASOLINE: detecting germline and somatic structural variants from long-reads data

**DOI:** 10.1101/2023.05.22.541558

**Authors:** Alberto Magi, Gianluca Mattei, Alessandra Mingrino, Chiara Caprioli, Chiara Ronchini, GianMaria Frigè, Roberto Semeraro, Davide Bolognini, Emanuela Colombo, Luca Mazzarella, Pier Giuseppe Pelicci

## Abstract

Long-read sequencing allows analyses of single nucleic-acid molecules and produces sequences in the order of tens to hundreds kilobases. Its application to whole-genome analyses allows identification of complex genomic structural-variants (SVs) with unprecedented resolution. SV identification, however, requires complex computational methods, based on either read-depth or intra- and inter-alignment signatures approaches, which are limited by size or type of SVs. Moreover, most currently available tools only detect germline variants, thus requiring separate computation of sample pairs for comparative analyses. To overcome these limits, we developed a novel tool (Germline And SOmatic structuraL varIants detectioN and gEnotyping; GASOLINE) that groups SV signatures using a sophisticated clustering procedure based on a modified reciprocal overlap criterion, and is designed to identify germline SVs, from single samples, and somatic SVs from paired test and control samples. GASOLINE is a collection of Perl, R and Fortran codes, it analyzes aligned data in BAM format and produces VCF files with statistically significant somatic SVs. Germline or somatic analysis of 30x sequencing coverage experiments requires 4-5 hours with 20 threads. GASOLINE outperformed currently available methods in the detection of both germline and somatic SVs in synthetic and real long-reads datasets. Notably, when applied on a pair of metastatic melanoma and matched-normal sample, GASOLINE identified 6 genuine somatic SVs that were missed using five different sequencing technologies and state-of-the art SV calling approaches. Thus, GASOLINE identifies germline and somatic SVs with unprecedented accuracy and resolution, outperforming currently available state-of-the-art WGS long-reads computational methods.

## Introduction

Structural variants (SVs) are genomic alterations typically defined (and somewhat arbitrarily) as DNA segments larger than 50 bp that can be deleted, duplicated, inserted, inverted or translocated compared to a reference genome. SVs are among the main sources of germline genomic variation in humans and can be associated with several diseases, including type I diabetes (8), cardiovascular disease (11), neurological disorders (22) and cancer (4). Moreover, somatic SVs acquired by cancer genomes are known drivers of carcinogenesis and their detection is essential for either diagnosis or treatment stratification in at least 30% of cancer patients (2).

In the past decade second-generation sequencing (SGS) technologies, based on high-throughput shortread generation (20), together with the development of powerful computational tools, have revolutionized our capability to study structural variations (SVs) of any size, from small insertions/deletions to large CNVs, with unprecedented accuracy in determining position and orientation (25). However, it is becoming apparent that the short reads (100-400 bp) generated by these platforms are insufficient to confidently detect variants larger than 50 bp, in particular those in the range [50, 500] bp (5).

The paste decade has seen the emergence of a third generation of sequencing technologies based on single-molecule real-time (Pacific Biosciences, PacBio) (9) and nanopore sequencing (Oxford Nanopore Technologies, ONT) (7), which interrogate single molecule of DNA and are capable to produce sequences much longer than those generated by SGS methods.

Recently, Chaisson et al. (5) and Huddleston et al. (13), by using deep PacBio sequencing data from two haploid human genomes, resolved the complete sequence of a large amount of SVs, showing that around 80% of these variants were missed by SGS data (with the greatest increase in sensitivity occurring for events smaller than 5 kb, in size). These two seminal papers demonstrated that the use of long read data can definitively enlarge the spectrum of detectable genetic variants, becoming the cutting-edge approach for the study of complex genomic structures.

Identification of SVs from long-reads data requires complex computational methods, which are based on either read-depth (depth of coverage, DOC) or intra- and inter-alignment SV signatures (split-read alignments) approaches (17). While the DOC approach is limited to the identification of large deletions and duplications (> 100 Kb) (19), gapped alignment methods allow detection of deletions, inversion and translocations of any size, with insertions and duplications limited by read length (17).

All split-read approaches consist of complex procedures in which the genomic coordinates of SV signatures are clustered on the basis of their reciprocal overlap to find groups of similar signatures that support the same SV. This step is critical for recovering all the signatures generated by each SV. Incomplete recovery can in fact lead to underestimation of the allelic fraction and genotyping errors. Signature clustering becomes even more critical when comparing datasets of matched test and control samples for somatic variant detection, where missing of some SV signatures in the control sample can lead to the calling false positive somatic variants.

At present, most of the currently available tools, such as SVIM (12), CuteSV (15) and Sniffles (24), only detect germline variants, using SV-signature clustering procedures that are based on classical reciprocal overlap. Remarkably, the standard approach for the detection of somatic variants consists in the application of these methods separately on each of the paired samples (test and control), discarding SVs with supporting reads in the control sample (26).

To overcome the limits of currently available methods, we developed GASOLINE (Germline And SO-matic structuraL varIants detectioN and gEnotyping) tool, a software that, exploiting a novel reciprocal overlap measure to cluster SV signatures is capable to detect germline SVs from the analysis of single samples as well as somatic SVs from the comparison of test and matched-normal samples. We tested our novel tool on synthetic and real long reads datasets and demonstrated its potential to detect germiline and somatic SVs with unprecedented accuracy and resolution.

## MATERIALS AND METHODS

When sequencing data are aligned to a reference genome, in principle, each SV subtype generates a typical pattern of mapped reads, named SV signature, which is then used to identify the underlying alteration. These signatures can be classified in two distinct categories: gapped alignment or split read alignments. Alignment algorithms use gap penalty (27) to account for genomic differences (alterations) occurring from insertions or deletions in the sequences. For example, a deletion, that is a lack of a sequence, generates a gap in the alignment of the read relative to a reference, while an insertion creates a gap in the alignment of the reference relative to the read. When genomic differences are too large and exceed gap penalty, mapping algorithms generate split-alignment, in which consecutive segments of the query sequence are mapped to disjoint regions in the reference and can have discordant orientation.

While gapped alignment generates two signature categories (insertions and deletions), the signatures arising from split reads can recognize, in principle every type of alteration: i) two consecutive segments mapping far apart with the same or opposite orientations define, respectively, deletion or inversion signatures; ii) overlapping coordinates define a duplication signature; iii) a read splitted in three segments, with the first and third segments closely mapped, define an insertion signature; iv) finally, when two consecutive segments mapped on different chromosomes define a translocation signature.

The first critical step of SVs detection consists in finding and grouping, all the signatures generated by each genomic alteration. From a computational point of view this step consists in clustering the genomic coordinates of SV signatures to find groups of intervals with large reciprocal overlap. Owing to the high error rate, the alignment of long read data can be very noisy and the genomic coordinates of SV signatures generated by the same event can be imprecise and have variances of tens of bp. In this situation, identifying and clustering SVs signatures generated by small events (50-500 bp) require small reciprocal overlap that takes into consideration noisy and imprecise alignments, while large SVs (tens or hundreds of kb), less affected by error rate, needs large reciprocal overlap to prevent the inclusion of signatures arising from other events.

Thus, the use of standard reciprocal overlap criteria can underestimate or completely miss signatures of small SVs (using large reciprocal overlap), or include wrong signatures in the identification of large SVs (using small reciprocal overlap may include signatures of other SVs).

### GASOLINE Germline

To overcome the limits of classical reciprocal overlap we introduced a novel normalized reciprocal overlap (NRO) criterion that allows grouping both small and large SV signatures with high accuracy, thus reducing the effect of imprecise alignment.

NRO mitigates the effect of imprecise alignment by taking into account both overlapping and non-overlapping regions of two SV signatures with the following formula:

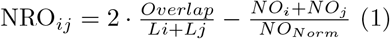

where *L*_*i*_ and *L*_*j*_ are the the size of the two signatures, *Overlap* and *NO* are the size of the overlapping and non-overlapping regions between the two signatures respectively and *NO*_*Norm*_ is a normalization factor. The first term represents the classical reciprocal overlap between two signatures, while the second term allows mitigation of the effect of imprecise alignments for small variants and prevents the inclusion of erroneous signatures in large SVs.

To identify germline SVs, GASOLINE takes as input the aligned data of a sample (in BAM format) and extract the genomic coordinates of gap- and split-alignment signatures (Figure 1.a1). SV signatures are then clustered with a sophisticated clustering procedure based on NRO measure, whose first step consists in calculating NRO for each pair of signatures (Figure 1.a2-a3). Once *NRO*_*ij*_ is calculated for all the signatures pairs [i,j] (Figure 1.b1), we perform interval clustering by using a graph-based approach. We first create an undirected graph by exploiting the NRO matrix as adjacency matrix, in which nodes are SV signatures and edges between two nodes *i* and *j* exist if *NRO*_*ij*_ *> Thr* (where *Thr* is a predefined reciprocal overlap threshold, Figure 1.b2). An edge between two nodes expresses the confidence of two signatures being generated by the same SV event (Figure 1.b3).

**Figure 1:**
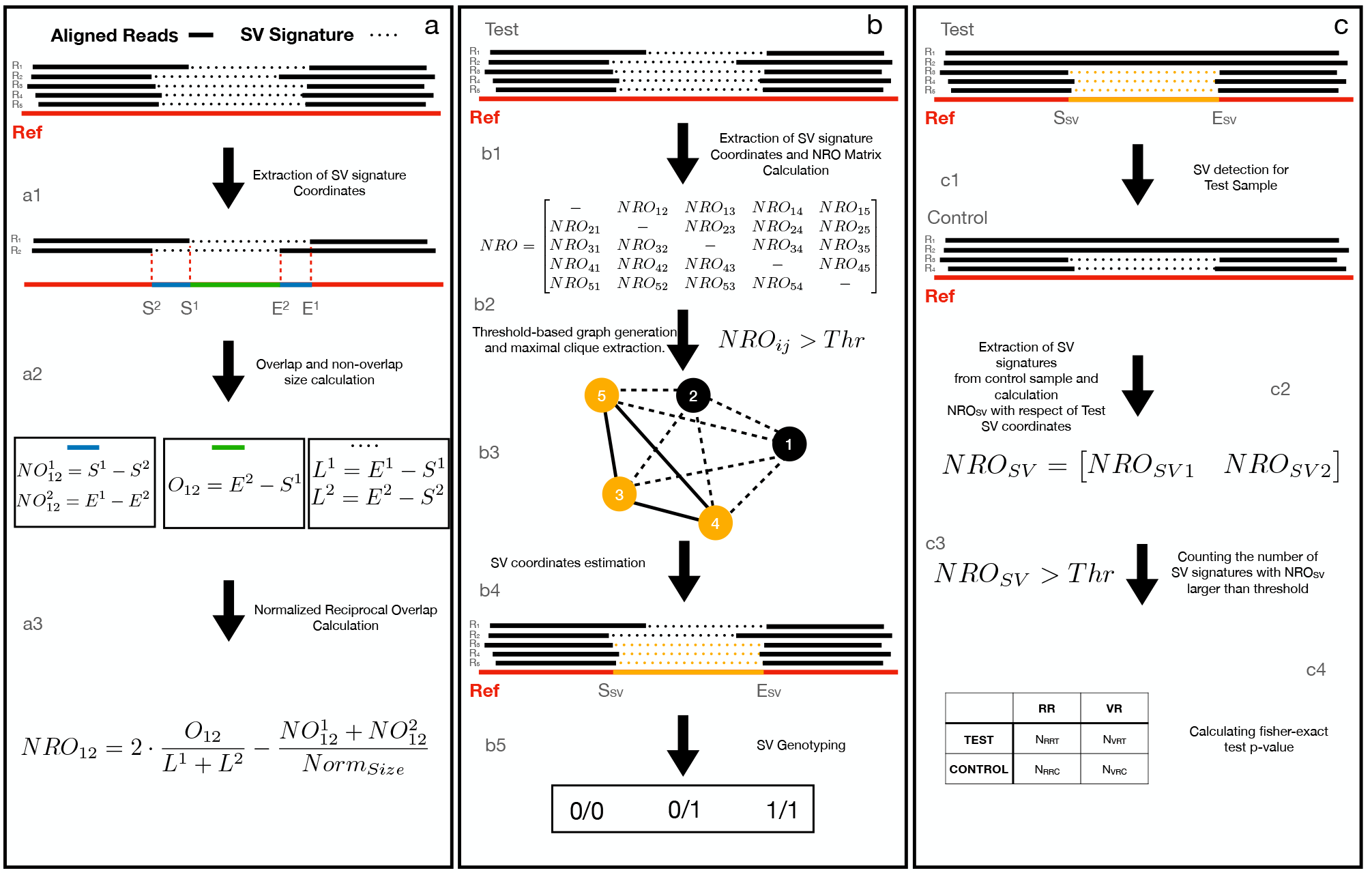
GASOLINE workflow. Panel (a) shows the steps to calculate NRO for a pair of SV signatures coordinates. Once the gap- and split-alignments coordinates (*S*_*i*_ and *E*_*i*_ for i,j=[1,2]) have been extracted from each read (a1), these are used to calculate the size of non-overlapping (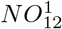 and 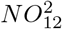, blue lines) and overlapping (*O*_12_) segments for each pair of signatures (a.2). 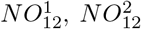, *O*_12_ and the total size of the two intervals (*L*^1^ and *L*^2^) are then used to calculate the *NRO*_12_ coefficient (a.3). In panel (b) is reported the workflow followed by GASOLINE for the detection of germline SVs in a sample. After signatures extraction, the tool calculates the *NRO*_*ij*_ between all the signature pairs and generates an NRO matrix (b.1) that is used as adjacency matrix to create an undirected graph by filtering out *NRO*_*ij*_ values smaller than a predefined threshold (continuous edges represent *NRO*_*ij*_ *> NRO*_*thr*_, while dotted edges *NRO*_*ij*_ *< NRO*_*thr*_, b2). The undirected graph is then analyzed with the Eppstein-Löffler-Strash algorithm to extract maximal cliques that represent clusters of SV signatures that can be assumed to be generated from the same SV event (b3). Next, all the SV signatures of a cluster are used to estimate the genomic coordinate (orange segment) of each SV event (b4). Finally, the number of SV signatures of a cluster and the total number of reads aligned in the breakpoints are used for genotyping with a maximum-likelihood Bayesian classification algorithm (b5). In panel (c) are reported the steps that GASOLINE follows for detecting somatic SVs. Somatic SVs are identified by comparing the SV signatures of a test (cancer) sample with a control (normal) sample. The SVs detected in the test sample (c1) are compared with the SVs signatures extracted from the control sample (c2) by calculating the NRO: SV signatures with a *NRO*_*SV*_ larger than a predefined threshold are considered to be generated from the SV event of the test sample (c3). Statistical significance of each somatic SV is calculated by applying the Fisher’s exact test on the contingency table of (c.4): *N*_*RRT*_ (number of reads without SV signatures in test sample), *N*_*VRT*_ (number of reads with the SV signatures in test sample), *N*_*RRC*_ (number of reads without SV signatures in control sample), *N*_*VRC*_ (number of reads with the SV signatures in control sample). SVs with a p-value smaller than a predefined significance threshold are considered somatic.

The undirected graph is then used to extract maximal cliques (groups of fully connected nodes) by using the Eppstein-Löffler-Strash algorithm (10). Maximal cliques represent groups of signatures that can be assumed to be generated from the same SV event (Figure 1.b3). SV signatures of each maximal clique are then used to estimate the genomic coordinates (start and end) of each SV event by calculating the median of all start and end coordinates (Figure 1.b4). For each cluster we then calculate different statistics including: cohesion score (the ratio between the numbers of links in the extended clique and the maximum numbers of link), mean mode and standard deviation of start and end coordinates. These statistics are then exploited to filter out low quality SV-signature clusters.

Finally, under the assumption of diploidy, each SV event is genotyped as reference, heterozygous, homozygous, by using the maximum-likelihood Bayesian classification algorithm as in (6) (Figure 1.b5).

### GASOLINE Somatic

The detection of somatic SVs in cancer genomes consists in the identification of SVs present in the cancer sample and absent in the patient-matched normal sample. GASOLINE identifies somatic SVs by first applying the germline detection module to the test data and then searching for overlapping SVs signatures in the matched normal sample (Figure 1.c).

The signature clusters identified in the test sample are then compared with the signatures extracted from the control sample, by calculating their NRO (Figure 1.c2). Signatures with a *NRO*_*SV*_ larger than a predefined threshold are considered to be generated from the SV event of the test sample (see Figure 1.c3).

Statistical significance of somatic SV is calculated by comparing the proportion between SV signatures and reference reads in the tumor and matched-normal samples with the Fisher’s exact test (contingency table of Figure 1.c4). SVs with a p-value smaller than a predefined significance threshold are considered somatic.

### PBSIM2

PBSIM2 simulates synthetic reads by randomly sampling from a reference sequence, adding errors with user defined distribution of substitutions, insertions and deletions and allowing to define read size distribution (mean and maximum size) and the desired sequencing coverage. PBSIM2 was exploited to simulate sequencing dataset that mimic the characteristics of long reads generated by ONT and PacBio platforms with total sequencing coverage from 5x to 30x (coverage=5x,10x, 15x, 20x, 25x and 30x). To study the performance of GASOLINE as a function of NRO values, we simulated all SV subtypes in homozygous and heterozygous state by applying PBSIM2 to reference sequences obtained by modifying a 5 Mb segment of chromosome 1 of the hg19 (1:5000001-10000000, see methods) with deleted, inserted, duplicated and inverted segments ranging from 50 bp to 5 kb (50 bp, 100 bp, 200 bp,… .). Translocations were simulated by applying PBSIM2 to two reference sequences obtained by modifying two 5 Mb segment of chromosome 1 of the hg19 (1:5000001-10000000 and 1:10000001-15000000, see methods). Simulated reads were then aligned to the 5 Mb reference genomes with minimap2 and NGMLR aligners. To compare the performance of GASOLINE with other three state of the art methods, we simulated inversions, duplications and translocations by combining SURVIVOR (https://github.com/fritzsedlazeck/SURVIVOR) and PBSIM2. SURVIVOR was used to generate a modified version of the Human Genome (hg19) with 3000 inversions, duplications and translocations ranging from 500 bp to 30 Kb. The modified and standard human genome was used to generate ONT and PacBio reads at 2.5x, 5x, 7.5x, 10x, 12.5x and 15x with PBSIM2. Reads from standard and modified genome were combined to obtain heterozygous SVs and aligned with minimap2 and NGMLR.

### Tools comparison

We downloaded the cuteSV tool (version 1.0.12) from https://github.com/tjiangHIT/cuteSV, Sniffles2 (version 2.0.7) from https://github.com/fritzsedlazeck/Sniffles and SVIM from https://github.com/eldariont/svim. For germline analyses, CuteSV was applied by using parameter settings suggested in the github page for ONT data (`--max cluster bias INS=100, --diff ratio merging INS=0.3, --max cluster bias DEL=100, --diff ratio merging DEL=0.3`) and for PacBio data (`--max cluster bias INS=100, --diff ratio merging INS=0.3, --max cluster bias DEL=200, --diff ratio merging DEL=0.5`), while Sniffles2 and SVIM were both run with default parameter settings.

Somatic analyses were performed by using each tool separately on paired samples (test and control) and discarding SVs with a supporting signature in the control sample. Supporting signatures in control samples were searched by using a reciprocal overlap larger than 0.1. GASOLINE was applied to all the datasets (PBSIM2 synthetic datasets, the NA24385 datasets, the COLO829 datasets and the synthetic somatic SVs generated with the NA24385 dataset), in ‘germline’ and ‘somatic’ mode with *NRO* = 0.8, *NO*_*Norm*_ = 1000. In all the analyses we performed, precision was calculated as the ratio between the number of correctly detected events (the intersection between the tool calls and the gold standard set calls) and the total number of events detected by each method, while recall was calculated as the ratio between the number of correctly detected events and the total number of events in the gold standard dataset (as in (18)). In both germline and somatic analyses each SV was considered a true positive if we found a reciprocal overlap larger than or equal to 50% with an SV of the validation set.

### NA24385 data

ONT and PacBio read data for the NA24385 individual of Ashkenazim ancestry was obtained from the GIAB ftp site https://ftp-trace.ncbi.nlm.nih.gov/giab/ftp/data/AshkenazimTrio/HG002\_NA24385\_son/. Reads in fastq (for ONT) or fasta (for PacBio) format were aligned against the human reference genome (hg19) by using minimap2 and NGMLR aligners, obtaining an average coverage of 64x for ONT and Xx for PacBio. To simulate 5, 10, 15, 20, 25 and 30x sequencing coverages, the original reads in fastq and fasta formats were downsampled by using the seqtk tool (https://github.com/lh3/seqtk) and then aligned to the human reference genome (hg19) with minimap2 and NGMLR. The GIAB consortium, by combining short-, long-, linked-read sequencing and optical mapping generated a high-quality callset of germline insertions and deletions. The NA24385 truth SV callset contains 12,745 SVs divided into 7,281 (6341 smaller than 1 kb and 787 in the range [1 kb, 5 kb]) insertions and 5,464 (4846 smaller than 1 kb and 462 in the range [1 kb, 5 kb]) deletions. The truth SV callset was downloaded at https://ftp-trace.ncbi.nlm.nih.gov/ReferenceSamples/giab/data/AshkenazimTrio/analysis/NIST\_SVs\_Integration\_v0.6/.

### COLO829 data

The COLO829 cancer cell-line from a metastatic cutaneous melanoma patient and the COLO829BL cellline from a lymphoblastoid line of the same patient were recently sequenced by Valle-Inclan et al. (26), by using five different technology platforms (Illumina HiSeq Xten, ONT, PacBio, 10x genomics and Bionano Genomics Saphyr optical mapping). By using state-of-the art SV calling approaches generated a somatic SV truth set comprising 68 high confidence calls: 38 deletions, 3 insertions, 7 duplications, 7 inversions and 13 translocations. The somatic SV truth set was downloaded from `10.5281/zenodo.3988185`. ONT, Illumina and PacBio WGS data were downloaded from EGA project PRJEB27698 (https://www.ebi.ac.uk/ena/browser/view/PRJEB27698). ONT and PacBio reads in fastq format were aligned against the human reference genome (hg19) by using minimap2 aligner, obtaining an average coverage of 60x for ONT and 50x for PacBio.

## Results

### GASOLINE and Germline SV detection

Many computational methods have been developed to detect SVs from different technologies and evaluation of their performance is a very challenging task, mainly due to the lack of gold-standard datasets including all subtypes of structural variation.

Thus, to assess the performance of GASOLINE in the detection and genotyping of germline SVs we first generated synthetic genomes with SVs of all subtypes and size by using the PBSIM2 software (21). PBSIM2 was exploited to simulate datasets mimicking the characteristics of long reads generated by either ONT or PacBio platforms, with average size of 10 Kb, a global error rate of 90% and a total sequencing coverage from 5x to 30x (coverage=5x,10x, 15x, 20x, 25x and 30x).We simulated all SV subtypes (deletions, insertions, duplications, inversions and translocations) in homozygous and heterozygous state and with size ranging from 50 bp to 5 kb (50, 100, 200, 300, 400, 500, 1000, 2000, 3000, 4000, 5000 bp). Simulated sequencing datasets were then aligned to the reference genome with minimap2 (16) and NGMLR (24) aligners (see methods).

To investigate the performance of our tool as a function of the NRO threshold, we applied it to the synthetic dataset using different parameter settings (NRO=[0.5, 0.6, 0.7, 0.8, 0.9]) and calculated precision and recall as in (18) (see methods). The results of these analyses (Supplemental Figures 1-3) demonstrated that NRO thresholds have little effect on the global performance of our tool, with NGMLR-generated data giving better results with low NRO thresholds (0.5-0.7), minimap2-generated data requiring instead higher values (0.8-0.9), particularly in the case of deletions.

Recently, the Genome in a Bottle (GIAB) Consortium, using a combination of short-, linked-, and long-read sequencing, as well as optical mapping, has characterized the genome of an individual of Ashkenazim ancestry (NA24385), thus generating gold-standard datasets for SVs (28). Although fundamental to test new technologies and algorithms, this dataset only contains high confidence sequence-resolved insertion and deletion calls > 50 base pairs (bp), and it does not enable performance assessment for inversions, duplications and translocations.

For this reason, we simulated inversions, duplications and translocation by using a computational strategy based on SURVIVOR (14) to modify the human reference genome and PBSIM2 to simulate ONT or PacBio reads (see Methods). Simulated reads were then aligned to the human reference genome with minimap2 (16) and NGMLR (24) aligners (see methods).

The synthetic dataset was then exploited to compare the performance of GASOLINE (using NRO=0.8) with those of other three state-of-the-art tools: Sniffles2 (24), CuteSV (15) and SVIM (12). The results reported in Figure 2.a-f and supplemental Figure 4 demonstrate that our method (and Sniffles2) obtained the best performance in the identification of simulated inversions, duplications and translocations, especially for low sequencing coverages (5-10x), thus demonstrating that our new NRO-based computational strategy is capable to group SV signatures with an high level of accuracy.

**Figure 2:**
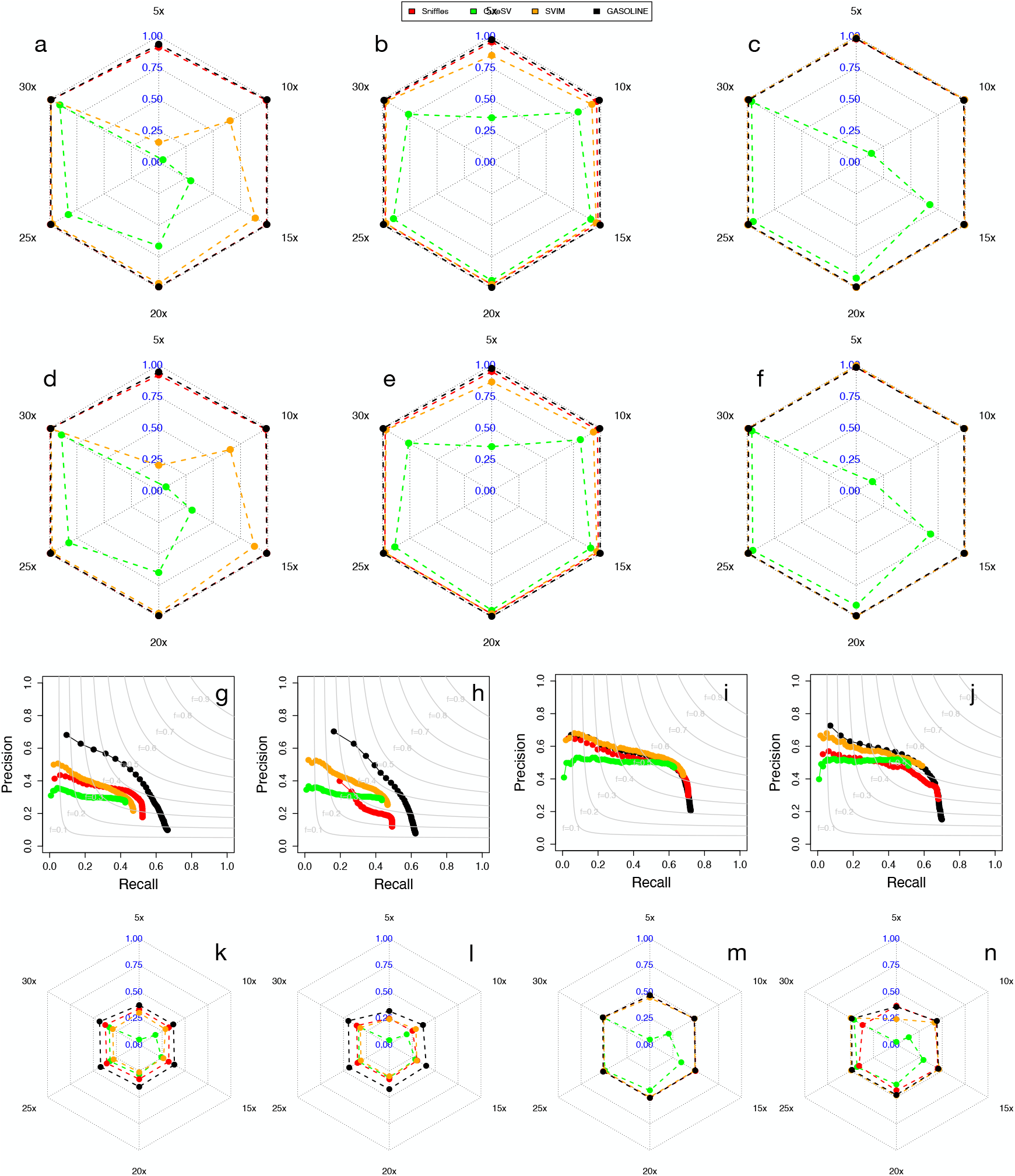
Global performance of GASOLINE and the other three tools in the detection of synthetic and real germline SVs. Panels a-f report the F1 score obtained by the four tools in the analysis of simulated inversions (a,d), duplications (b,e) and translocations (c, f). Results are reported for ONT (a-c) and PacBio (d-e) synthetic reads aligned with minimap2. Panels g-j report precision and recall the four tools in the detection of small SVs (g-h) and large SVs (i-j) with ONT (g, i) and PacBio (h, j) data. The curves in panels (g-j) were obtained by ordering all the SVs as a function of number of supporting reads and calculating precision and recall including SVs with decreasing number of reads. Panels k-n show F1 score obtained by the four tools in the detection of small (k-l) and large SVs (m-n) with ONT (k, m) and PacBio (l, n) datasets at different sequencing coverages.

Remarkably, sequencing coverage has little effects on the global performance of our tool for both alignment algorithms: as previously reported by (3) a sequencing coverage higher than 15x is sufficient for detecting all subtypes of SVs and its increase does not lead to significant improvements.

To evaluate the accuracy of GASOLINE in the detection of real germline insertions and deletions we applied it to the publicly available ONT and PacBio NA24385 datasets generated by the GIAB consortium (see Methods) and compared its performance with the three aforementioned tools. To assess the performance of the four tools in detecting SVs at different sequencing coverages, we downsampled ONT and PacBio datasets to simulate 5, 10, 15, 20, 25 and 30x coverages and performed alignments with both minimap2 and NGMLR.

To compare the capability of the four tools to cluster SV signatures, we calculated precision and recall as a function of numbers of reads supporting each SV event, separately for large and small SVs (see methods). Figure 2.g-j and supplemental Figures 5-16 show that for large insertions and deletions (> 500*bp*) all the tools performed very similarly. For small SVs (< 500*bp*), instead, GASOLINE showed the superiority of its NRO-based clustering procedure in grouping true SV signatures from long reads noisy alignments, obtaining the highest precision at the same level of recall for all sequencing coverages (Supplemental Figures), sequencing data (PacBio and ONT) and aligners (NGMLR and minimap2). These analyses also showed that NGMLR alignment gave better results in the detection of large SVs, while minimap2 data better detected small SVs.

Finally, we calculated precision and recall for all SVs genotyped as heterozygous (0/1) or homozygous (1/1) by the four tools and we found that while for large insertions and deletions (> 500*bp*) all tools obtained very similar results, for small SVs (< 500*bp*) GASOLINE obtained the highest F1 measure for all sequencing coverages and technologies (Figure 2.k-n).

### Somatic SV detection on simulated and real data

As for germilne variants, the validation of computational methods for somatic SVs detection is challenged by the lack of high-quality gold standard datasets enabling benchmarking and comparison of bioinformatic analysis pipelines, especially for tools exploiting long-read datasets and capable to identify small and complex variants previously unseen by short-reads WGS experiments.

Recently, Valle-Inclan et al. (26), generated a comprehensive set of true somatic SVs (comprising all SV types) of the melanoma COLO829 cell lines by using four different sequencing technologies (Illumina HiSeq, ONT, PacBio and 10x Genomics) combined with extensive experimental validation (see Methods). Despite the great utility of such a gold reference dataset, its application for benchmarking purposes on long read data is limited by the small number of insertions and by the size distribution of all the SVs that are mainly larger than 50 kb not allowing to evaluate the performance of algorithms in the detection of small SVs ([50,500] bp).

For these reasons, to assess the accuracy of GASOLINE in the identification of small somatic insertions and deletions we simulated somatic SVs of different sizes by using the PacBio and ONT WGS NA24385 dataset generated by the GIAB consortium (see Methods). We selected 330 heterozygous SVs (169 insertions and 161 deletions from the high-confidence callsets generated by the GIAB consortium) with size distribution in the range [50*bp*, 5*kb*] (with 230 SVs smaller than 1 kb). Using custom scripts, we first removed all reads containing signatures of the 330 SVs, we randomly splitted the remaining reads in two sets and finally, we added all the reads containing SV signatures to one of the splitted sets. With this workflow, for both PacBio and ONT data we obtained a *∼* 30*x* sequencing experiment with reads containing the 330 selected SVs (test), and a *∼* 30*x* control experiment without the 330 SVs. To evaluate detection accuracy at different sequencing coverages we downsampled read datasets to obtain 5x, 10x, 15x, 20x and 25x. Raw reads were aligned using either minimap2 or NGMLR. We then applied the somatic module of GASOLINE to the simulated datasets and we compared its performance to the other three tools (see methods).

GASOLINE obtained the best F-measure for all sequencing coverages of both PacBio and ONT datasets aligned with minimap2 or NGMLR (Figure 3.a-d and supplemental figure 17-28), especially in the detection of small SVs ([50 *−* 500]). In contrast, the other three tools identified a large number of false positive somatic SVs resulting in very low levels of precision. These analyses also showed that low sequencing coverages (*≤* 15*x*) yield very poor performance and that, similarly to germline SVs, the best F1-measure is obtained with at least 20x coverage, for both PacBio and ONT data. Notably, ONT data outperformed PacBio in all our analyses, while the minimap2 approach showed higher precision and recall than obtained with NGMLR.

**Figure 3:**
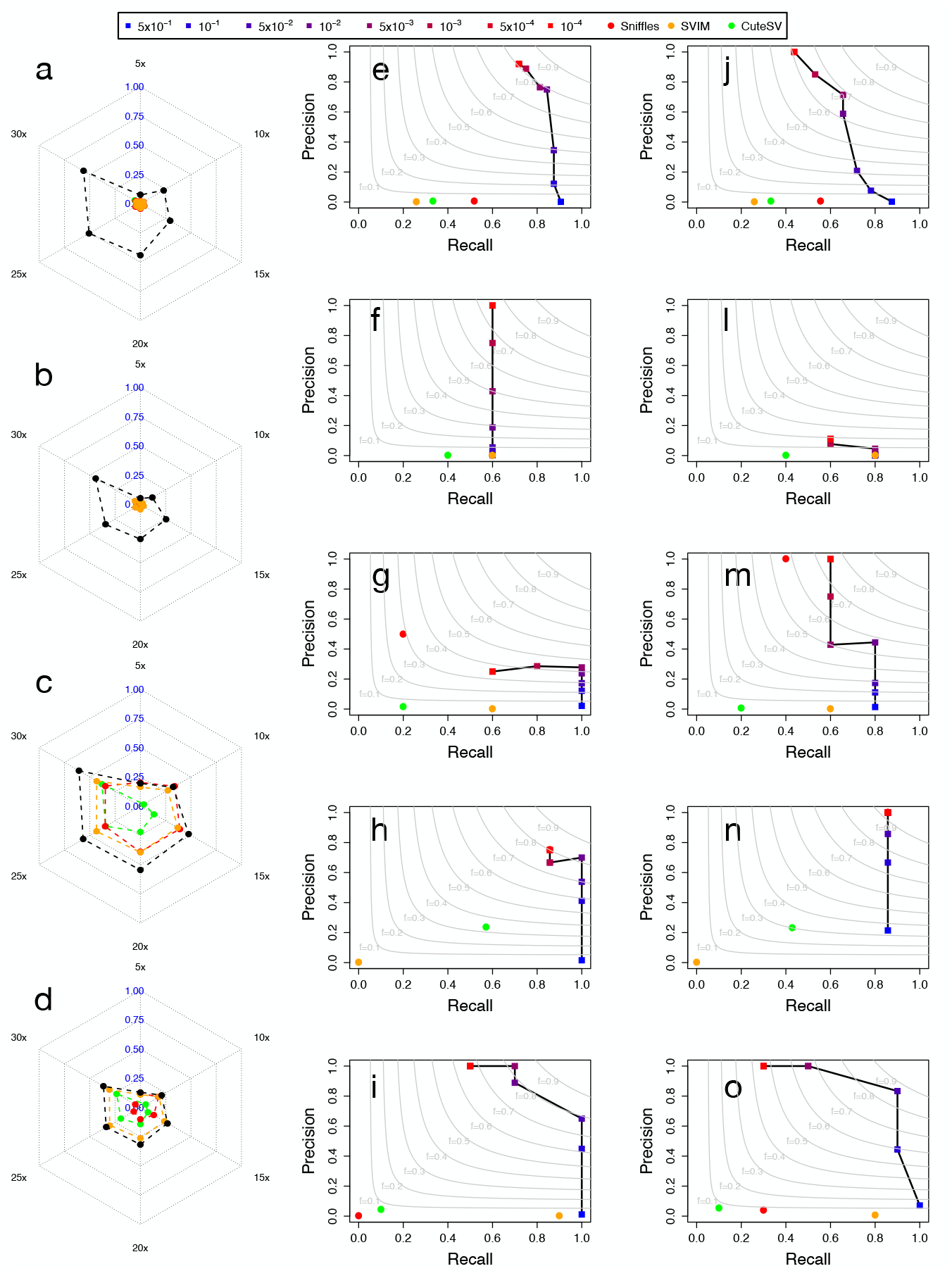
Performance of GASOLINE on the detection of somatic SVs. Panels a-d show F1 score obtained by the four tools in the detection of simulated small (a-b) and large somatic SVs (c-d) with ONT (a, c) and PacBio (b, d) datasets at different sequencing coverage. Results for GASOLINE Panels (e-o) report the precision-recall obtained by GASOLINE and the other three tools in the detection of deletions (e,j), insertions (f, l), duplications (g, m), inversions (h, n) and translocations (i, o) of the Valle-Inclan et al. true-set for the COLO829 cell lines sequenced with ONT (e-i) and PacBio (j-o) technologies. The results for GASOLINE were reported for different somatic p-value thresholds (5×10^*−*1^, 1×10^*−*1^, 5×10^*−*2^, 1×10^*−*2^, 5×10^*−*3^, 1×10^*−*3^, 5×10^*−*4^, 1×10^*−*4^). All the results reported in the panels are based on mimimap2 alignment data.

We next tested the capability of our tool in detecting all SVs subtypes, applying it to the ONT and PacBio data of the COLO829 cell lines and comparing its performance with those of the other three tools by using the Valle-Inclan et al. (26) true-set as benchmark (see Methods). As with the simulated datasets, the three state of the art tools identified a large number of false positive somatic events that generated very low levels of precision. On the other hands, GASOLINE, filtering out SVs on the basis of somatic p-values, was capable to drastically increase precision (removing a large fraction of false positive calls) at the expense of a minimal decrease in recall (removal of true positive calls). For all SV subtypes, with the exception of insertions (unfortunately the gold standard true dataset only contains three somatic insertions), our method was capable to identify between 80% and 100% of the Valle-Inclan et al. true-set with precision in the order of 60-80%, outperforming the other three state of the art methods (Figure 3.e-o) and demonstrating that GASOLINE can expand our potential to study somatic alterations in cancer.

### GASOLINE tool

GASOLINE is a collection of Perl, R and Fortran codes for the detection of somatic SVs from long read sequencing data. It takes as input two BAM files from a pair of test and matched normal samples and gives as output a VCF file (version 4.2) with statistically significant somatic SVs.

GASOLINE analyzes aligned data in BAM format and extracts the genomic coordinates of discordant alignments of a read with respect to the reference genome (SV signatures). The parsing module of our tool searches for two types of signatures: gapped alignments (in the CIGAR strings) and split alignments (primary and supplementary alignment of a read).

At present the signature extraction module is written in Perl and takes around three hours for parsing a ONT or PacBio WGS at 30x of sequencing coverage. After the parsing step, GASOLINE can be run in ‘germline’ or ‘somatic’ mode. In ‘germline’ mode, SV signatures are grouped with the NRO-based clustering and then genotyped. In ‘somatic’ mode SV signatures from a test cancer sample are clustered and then compared with those of matching control sample to calculate somatic p-value with Fisher’s exact test.

Both ‘germline’ and ‘somatic’ modules can be ran in multicores: the ‘germline’ module takes around one hour to genotype a 30x bam file (with 20 threads), while the ‘somatic’ module takes around two hours two compare the SV signatures of two 30x bam files (with 20 threads).

GASOLINE can run on any UNIX system (desktops and workstations). The GASOLINE tool is freely available at https://sourceforge.net/projects/gasoline/.

## Discussion and Conclusion

Long-read sequencing technologies are revolutionizing our capability og identifying and resolve the structure of complex SVs with an unprecedented accuracy and resolution. However, currently available tools for long-read analyses are based on computational procedures that limit the detection of small SVs (< 500 bp). Notably, none of them is properly devised for the identification of somatic alterations.

In order to overcome the limits of currently available methods, we developed GASOLINE, the first computational approach that is capable of detecting germline and somatic SVs from long reads sequencing datasets. GASOLINE is based on a novel reciprocal overlap (NRO) criterion that allows to group both small and large SV signatures with high accuracy, thus reducing the effect of imprecise alignment and allowing the identification of both small (< 500 bp) and large (> 500 bp) SVs with the same accuracy. Analyses of synthetic and real long-read datasets demonstrated that our NRO-based clustering algorithm clearly outperform the other state of the art method in the detection of germline alterations, especially SVs smaller than 500 bp.

The most important novelty and uniqueness of GASOLINE lies in its capability to compare a test and a matched normal sample to identify somatic alterations. At present, the standard approach for the detection of somatic variants consists in applying these methods separately on paired samples (test and control) discarding SVs with a supporting read in the control sample. GASOLINE directly compares the SV signatures found in test and control samples and then it calculates somatic statistical significance with Fisher’s exact test.

As for germline variants we tested the performance of our tool in the detection somatic variants, using both simulated and real long-read cancer datasets. In synthetic datasets, as for germline variants, our tool demonstrated its superiority in the detection of small SVs for all sequencing coverages we simulated. In particular, the somatic p-values calculated by GASOLINE are a useful instrument to increase precision (removing a large fraction of false positive calls) at the expense of a minimal decrease in recall (removal of true positive calls).

When applied on a pair of metastatic cutaneous melanoma (COLO829) and matched normal sample, GASOLINE outperformed the other three tools in the detection of all SV subtypes.

At present, the speed of GASOLINE is, however, still slower than the other of state-of-the-art approaches. At present, the germline or somatic analysis of a 30x coverage sequencing experiments requires 4-5 hours with 20 threads. This is mainly due to the SV signature extraction module of our method that is implemented in perl. We are planning to implement the parsing module in *c* + +, to obtain computational speed comparable to that of currently available state of the art SV callers.

Regardless, the results obtained in all of the comparative analyses we performed highlighted the versatility of our software and its ability to overcome the limitations and drawbacks of currently available state-of-the-art tools, thus making GASOLINE a suitable tool for the investigation of SVs in population as well as in cancer studies.

## Author contributions statement

Alb.M. conceived and implemented GASOLINE. Alb.M., G.M. R.S. and D.B. performed all bioinformatic analyses. Alb.M, P.G.P., G.M., L.M. and C.C. contributed to results interpretation. Alb.M. and P.G.P. wrote the manuscript. All the authors reviewed and edited the manuscript.

## References

[1] Belyeu JR, Chowdhury M, Brown J, Pedersen BS, Cormier MJ, Quinlan AR, Layer RM. Samplot: a platform for structural variant visual validation and automated filtering. Genome Biol. 2021 May 25;22(1):161.

[2] van Belzen IAEM, Schönhuth A, Kemmeren P, Hehir-Kwa JY. Structural variant detection in cancer genomes: computational challenges and perspectives for precision oncology. NPJ Precis Oncol. 2021 Mar 2;5(1):15.

[3] Bolognini D, Magi A. Evaluation of Germline Structural Variant Calling Methods for Nanopore Sequencing Data. Front Genet. 2021 Nov 18;12:761791.

[4] Campbell PJ et al. Identification of somatically acquired rearrangements in cancer using genome-wide massively parallel paired-end sequencing. Nat Genet. 2008 Jun;40(6):722–9.

[5] Chaisson MJ et al. Resolving the complexity of the human genome using single-molecule sequencing. Nature. 2015 Jan 29;517(7536):608–11.

[6] Chiang C, Layer RM, Faust GG, Lindberg MR, Rose DB, Garrison EP, Marth GT, Quinlan AR, Hall IM. SpeedSeq: ultra-fast personal genome analysis and interpretation. Nat Methods. 2015 Oct;12(10):966–8.

[7] Clarke J et al. Continuous base identification for single-molecule nanopore DNA sequencing. Nat Nanotechnol 2009;4(4):265?70.

[8] Craddock N, Hurles ME, Cardin N, Pearson RD et al. Genome-wide association study of CNVs in 16,000 cases of eight common diseases and 3,000 shared controls. Nature. 2010 Apr 1;464(7289):713–20.

[9] Eid J et al. Real-time DNA sequencing from single polymerase molecules. Science. 2009 Jan 2;323(5910):133–8.

[10] Eppstein D, Löffler M and Strash D. Listing All Maximal Cliques in Sparse Graphs in Near-optimal Time. arXiv:1006.5440

[11] Fahed, A. C., Gelb, B. D., Seidman, J. G., Seidman, C. E. (2013). Genetics of congenital heart disease: the glass half empty. Circulation research, 112(4), 707–720.

[12] Heller D, Vingron M. SVIM: structural variant identification using mapped long reads. Bioinformatics. 2019 Sep 1;35(17):2907–2915.

[13] Huddleston J et al. Discovery and genotyping of structural variation from long-read haploid genome sequence data. Genome Res. 2017 May;27(5):677–685.

[14] Jeffares DC, Jolly C, Hoti M, et al. Transient structural variations have strong effects on quantitative traits and reproductive isolation in fission yeast. Nat Commun. 2017;8:14061.

[15] Jiang T, Liu Y, Jiang Y, Li J, Gao Y, Cui Z, Liu Y, Liu B, Wang Y. Long-read-based human genomic structural variation detection with cuteSV. Genome Biol. 2020 Aug 3;21(1):189.

[16] Li H. Minimap2: pairwise alignment for nucleotide sequences. Bioinformatics. 2018 Sep 15;34(18):3094–3100.

[17] Mahmoud M, Gobet N, Cruz-Dávalos DI, Mounier N, Dessimoz C, Sedlazeck FJ. Structural variant calling: the long and the short of it. Genome Biol. 2019 Nov 20;20(1):246.

[18] Magi A et al. EXCAVATOR: detecting copy number variants from whole-exome sequencing data. Genome Biol. 2013;14(10):R120.

[19] Magi A et al. Nano-GLADIATOR: real-time detection of copy number alterations from nanopore sequencing data. Bioinformatics. 2019 Nov 1;35(21):4213–4221.

[20] Metzker ML. Sequencing technologies - the next generation. Nat Rev Genet. 2010 Jan;11(1):31–46.

[21] Ono Y, Asai K, Hamada M. PBSIM2: a simulator for long-read sequencers with a novel generative model of quality scores. Bioinformatics. 2021 May 5;37(5):589–595.

[22] Pippucci T et al. Epilepsy with auditory features: A heterogeneous clinico-molecular disease. Neurol Genet. 2015 May 14;1(1):e5.

[23] Robinson JT, Thorvaldsdóttir H, Winckler W, Guttman M, Lander ES, Getz G, Mesirov JP. Integrative genomics viewer. Nat Biotechnol. 2011 Jan;29(1):24–6.

[24] Sedlazeck FJ, Rescheneder P, Smolka M, Fang H, Nattestad M, von Haeseler A, Schatz MC. Accurate detection of complex structural variations using single-molecule sequencing. Nat Methods. 2018 Jun;15(6):461–468.

[25] Tattini L, D’Aurizio R, Magi A. Detection of Genomic Structural Variants from Next-Generation Sequencing Data. Front Bioeng Biotechnol. 2015 Jun 25;3:92.

[26] Valle-Inclan JE, Besselink NJ, de Bruijn E, Cameron DL, Ebler J, Kutzera J, Van Lieshout S, Marschall T, Nelen M, Pang AW, Priestley P. A multi-platform reference for somatic structural variation detection. bioRxiv 2020.10.15.340497; doi: https://doi.org/10.1101/2020.10.15.340497

[27] Vingron M, Waterman MS. Sequence alignment and penalty choice. Review of concepts, case studies and implications. J Mol Biol. 1994 Jan 7;235(1):1–12.

[28] Zook JM et al. A robust benchmark for detection of germline large deletions and insertions. Nat Biotechnol. 2020 Nov;38(11):1347–1355.

