## Supplemental Material for "GASOLINE: detecting germline and somatic structural variants from long-reads data"

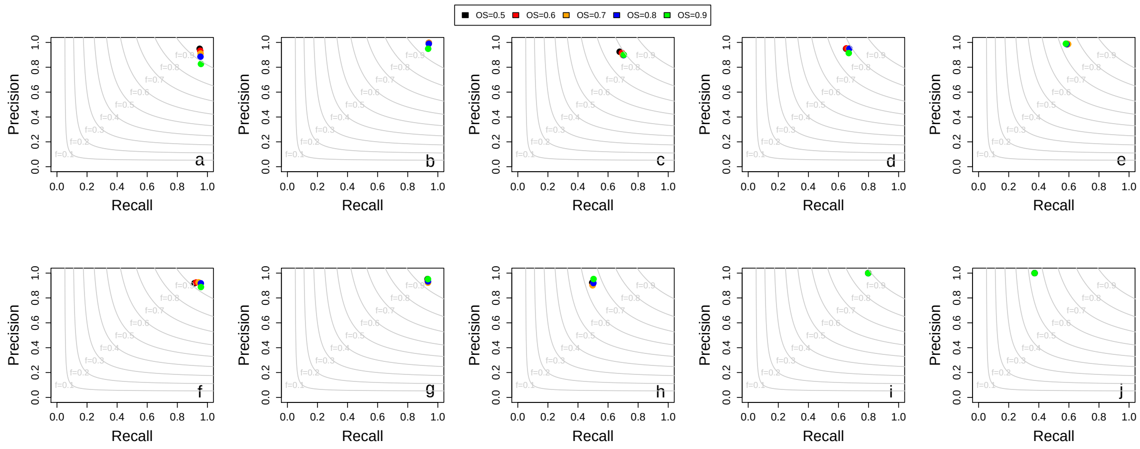


Figure 1: Global performance of GASOLINE and the other three tools on synthetic data. Panels a-e report the precision-recall plots for synthetic reads aligned with NGMLR, while panels f-j for synthetic reads aligned with minimap2. Panels a and f reports the performance of the four methods in the detection of synthetic deletions, panels b and g for synthetic insertions, panels c and h for duplications, d and I for inversions and e and j for translocations.

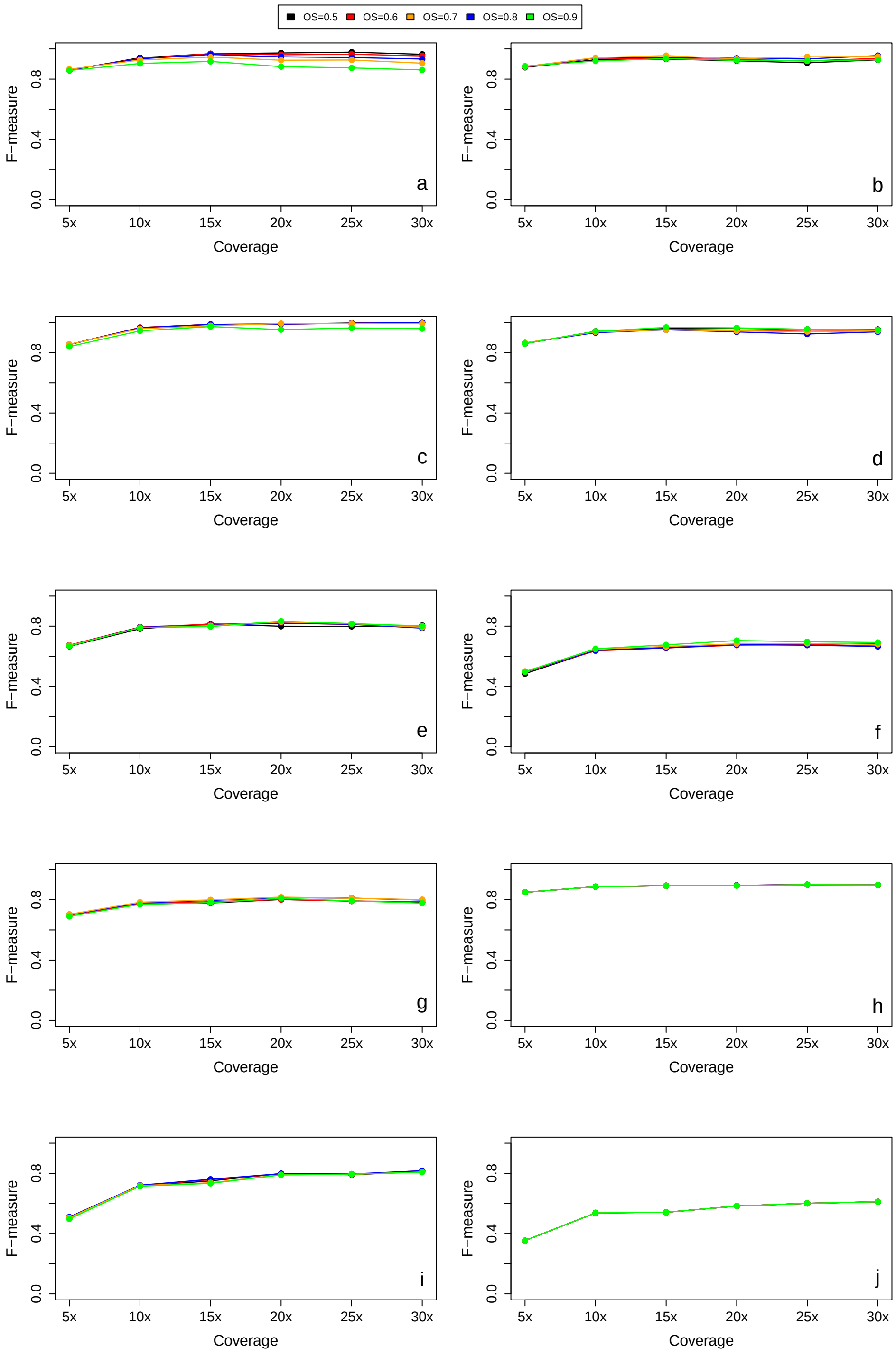


Figure 2: Performance of GASOLINE with different NRO values on synthetic data as a function of sequencing coverage. Figure reports the armonic mean of precision and recall (F-measure) obtained by GASOLINE in the detection of deletions (panels a and b), insertions (panels c and d), duplications (panels e and f), inversions (panels g and h) and translocations (panels i and j) from simulated data with different sequencing covarages (5x, 10x, 15x, 20x, 25x, 30x). Result are reported for sequencing data aligned with minimap2 (panels a, c, e, g, i) and NGMLR (panels b, d, f, h, j). GASOLINE was tested for different NRO values thresholds (NRO=0.5, NRO=0.6, NRO=0.7, NRO=0.8, NRO=0.9).

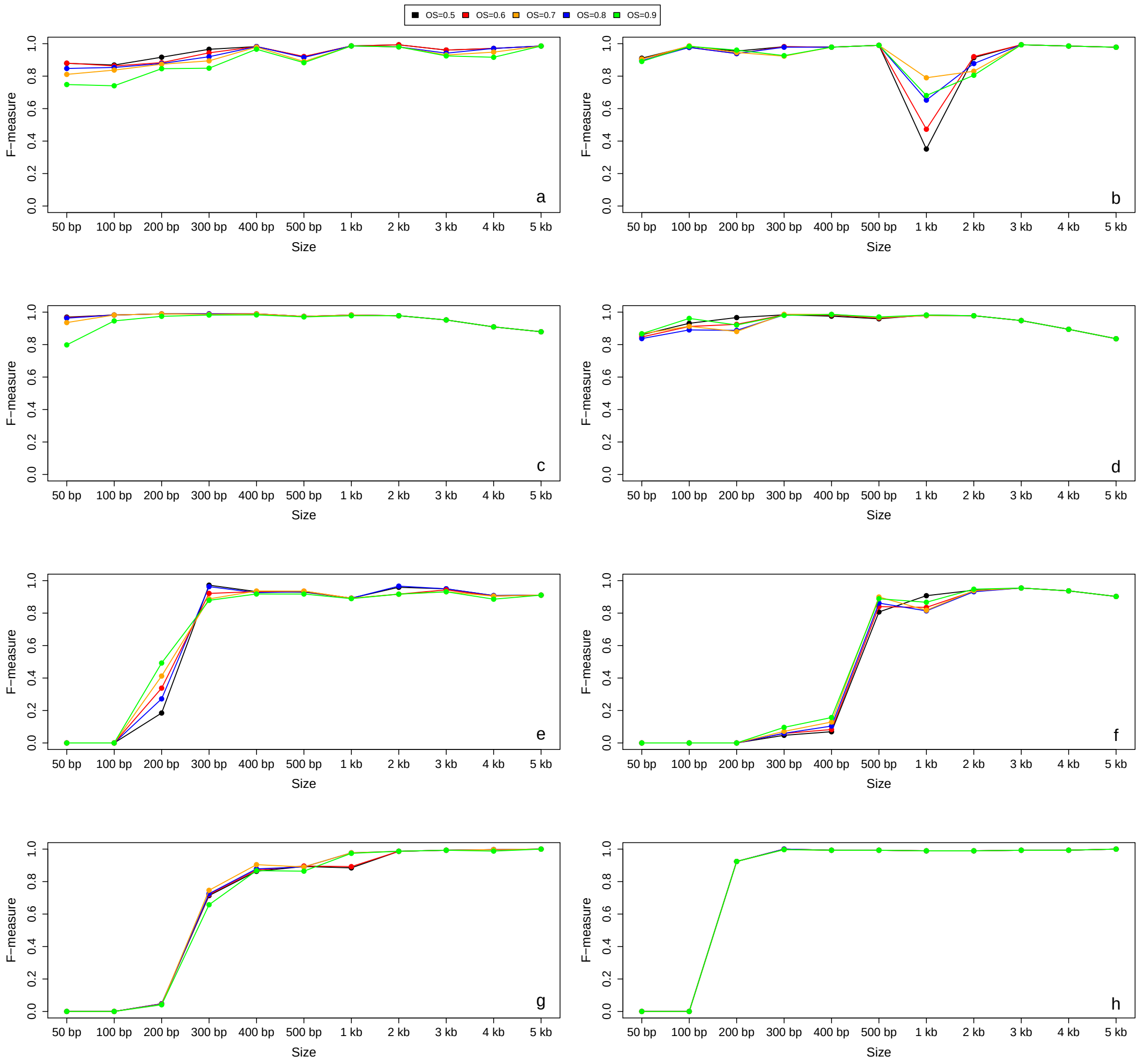


Figure 3: Performance of GASOLINE with different NRO values on synthetic data as a function of SV size. Figure reports the armonic mean of precision and recall (F-measure) obtained by GASOLINE in the detection of deletions (panels a and b), insertions (panels c and d), duplications (panels e and f), inversions (panels g and h) and translocations (panels i and j) of different size (50 bp, 100 bp, 200 bp, 300 bo, 400 bp, 500 bp, 1 kb, 2 kb, 3 kb, 4 kb, 5 kb). Result are reported for sequencing data aligned with minimap2 (panels a, c, e, g, i) and NGMLR (panels b, d, f, h, j). GASOLINE was tested for different NRO values thresholds (NRO=0.5, NRO=0.6, NRO=0.7, NRO=0.8, NRO=0.9).

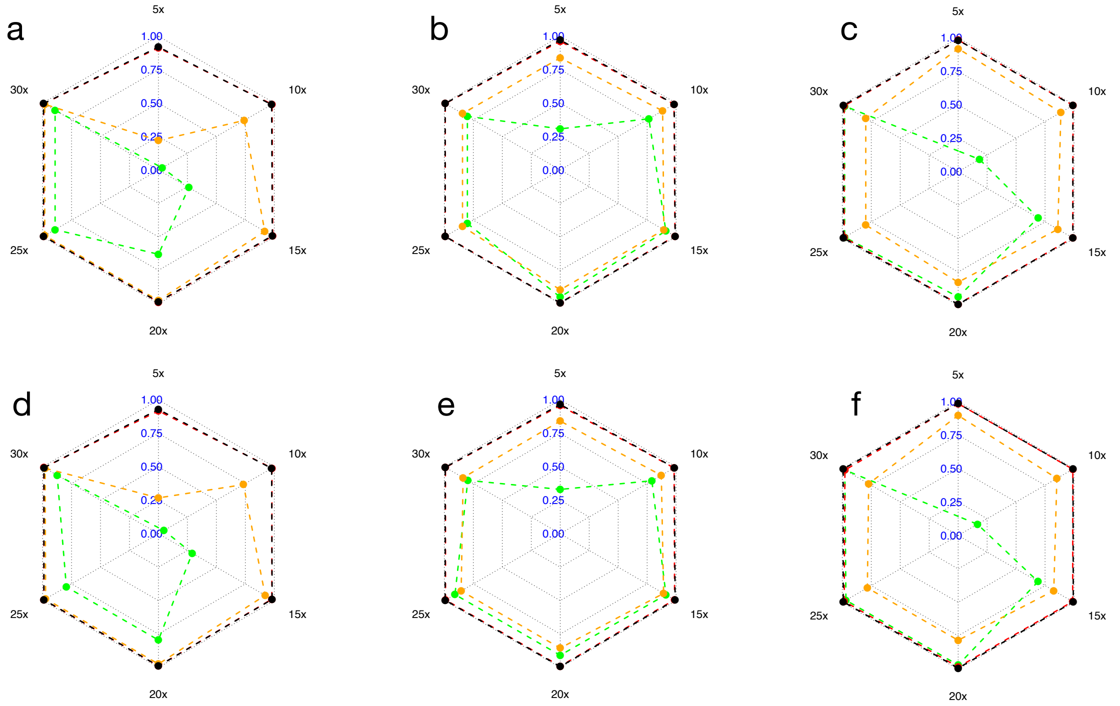


Figure 4: Panels a-f report the F1 score obtained by the four tools in the analysis of simulated inversions (a,d), duplications (b,e) and translocations (c, f). Results are reported for ONT (a-c) and PacBio (d-e) synthetic reads aligned with NGMLR.

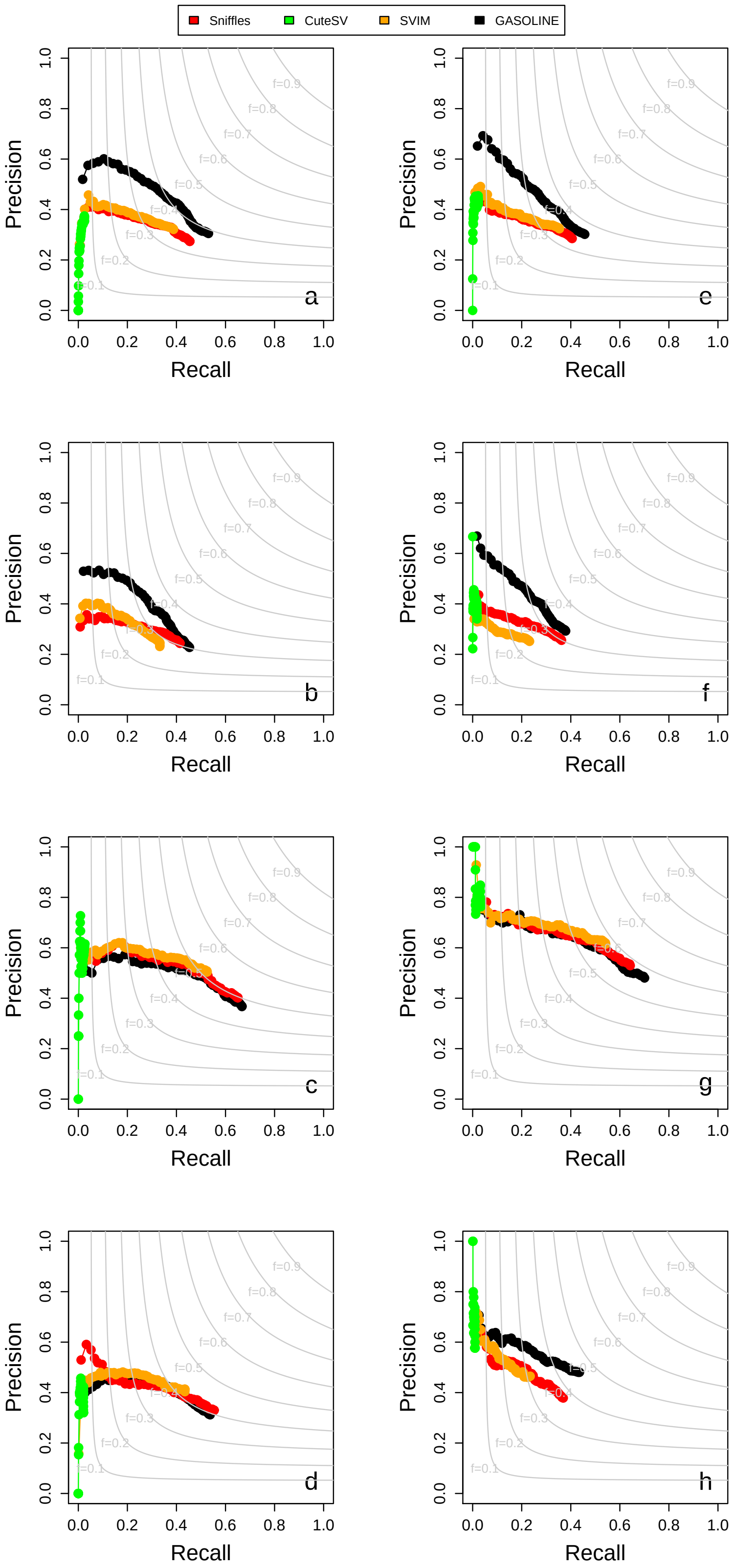


Figure 5: Precision-Recall curves of GASOLINE and the other three tools as a function of number of supporting reads for NA24385 ONT data downsampled at 5x. Panels (a-d) show the results for minimap2 alignment and (e-h) for NGMLR. Panels (a, e) for small deletions (< 500 bp), (b, f) for small insertions (< 500 bp), (a, e) for large deletions (> 500 bp), (a, e) for large insertions (> 500 bp). The curves in panels were obtained by ordering all the SVs as a function of number of supporting reads and calculating precision and recall including SVs with decreasing number of reads.

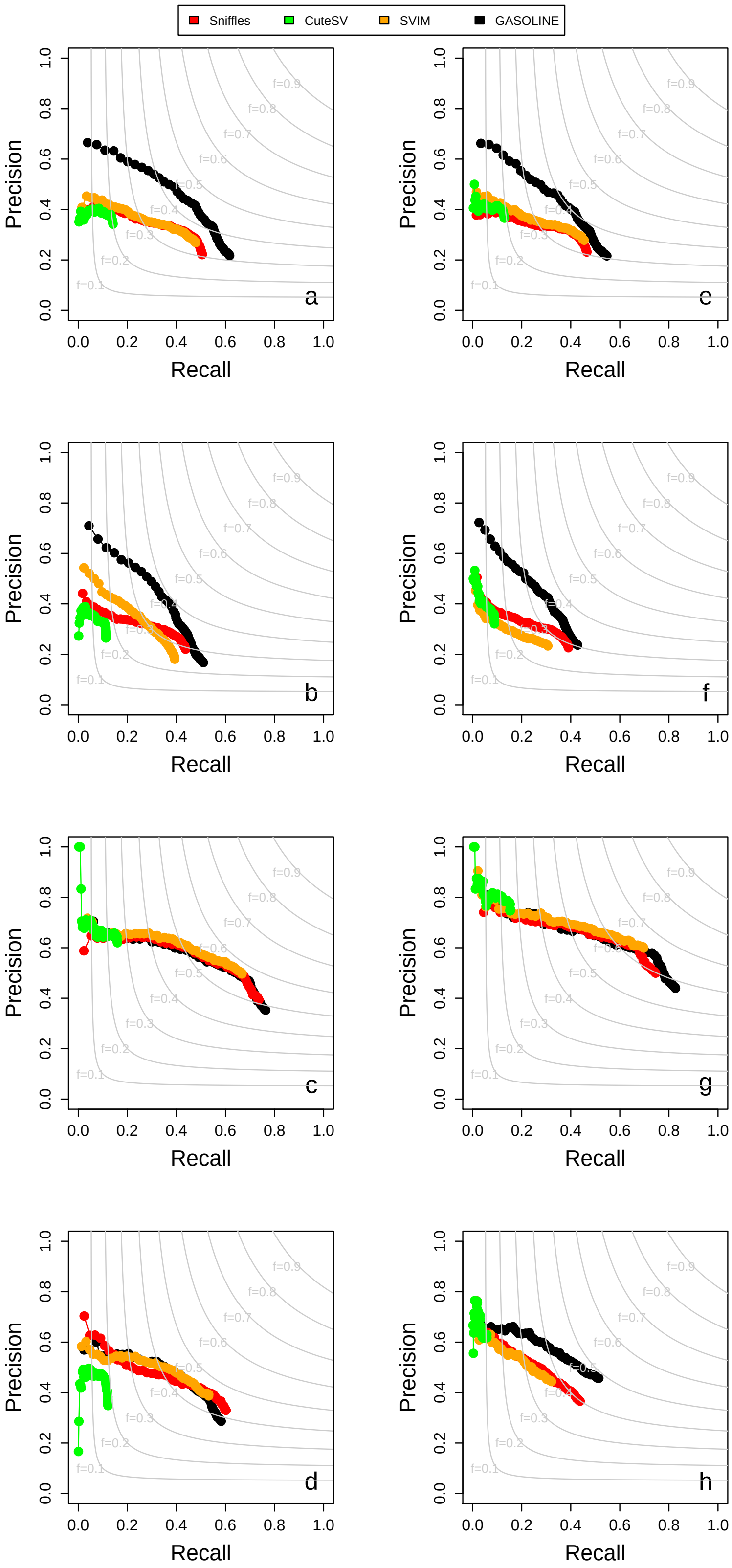


Figure 6: Precision-Recall curves of GASOLINE and the other three tools as a function of number of supporting reads for NA24385 ONT data downsampled at 10x. Panels (a-d) show the results for minimap2 alignment and (e-h) for NGMLR. Panels (a, e) for small deletions (< 500 bp), (b, f) for small insertions (< 500 bp), (a, e) for large deletions (> 500 bp), (a, e) for large insertions (> 500 bp). The curves in panels were obtained by ordering all the SVs as a function of number of supporting reads and calculating precision and recall including SVs with decreasing number of reads.

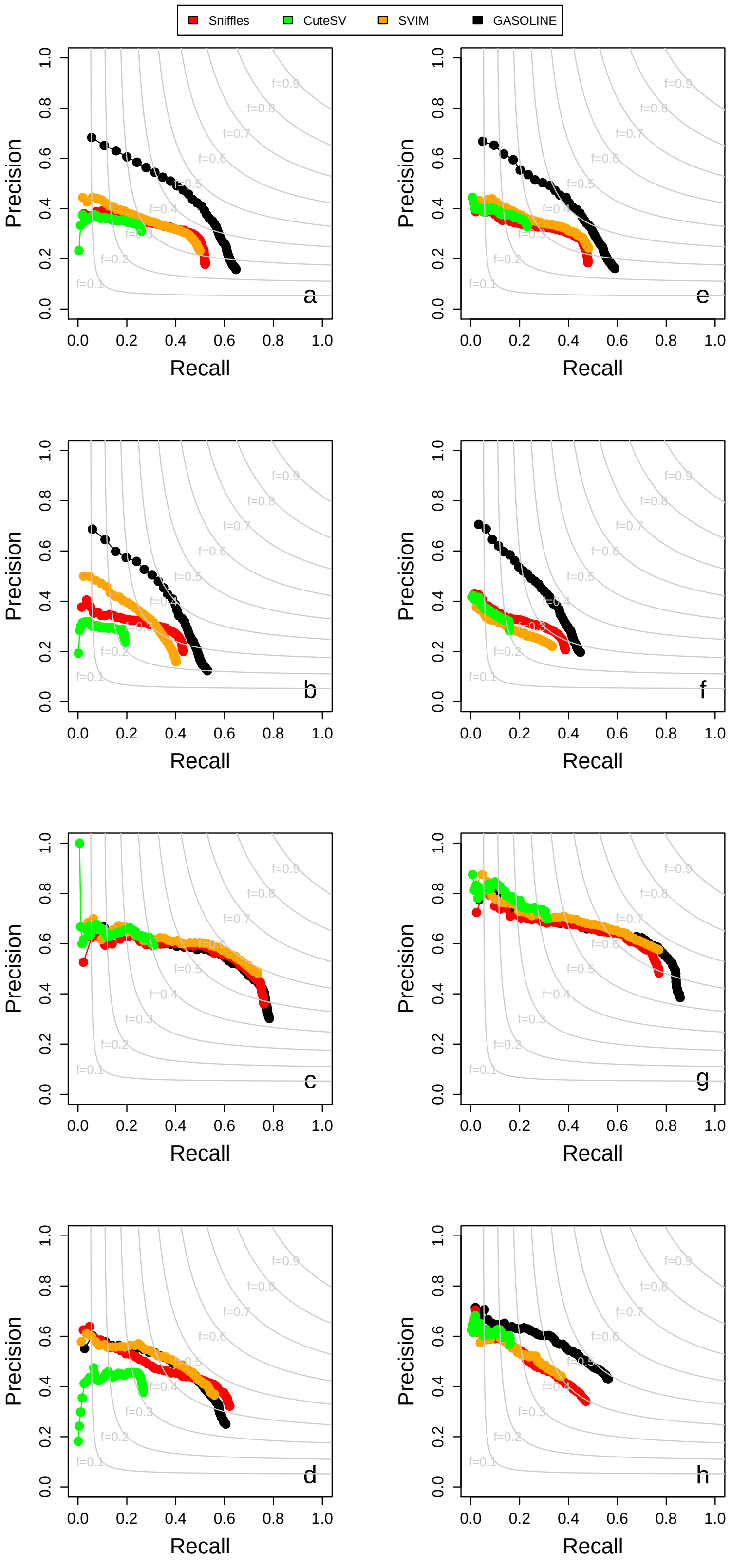


Figure 7: Precision-Recall curves of GASOLINE and the other three tools as a function of number of supporting reads for NA24385 ONT data downsampled at 15x. Panels (a-d) show the results for minimap2 alignment and (e-h) for NGMLR. Panels (a, e) for small deletions (< 500 bp), (b, f) for small insertions (< 500 bp), (a, e) for large deletions (> 500 bp), (a, e) for large insertions (> 500 bp). The curves in panels were obtained by ordering all the SVs as a function of number of supporting reads and calculating precision and recall including SVs with decreasing number of reads.

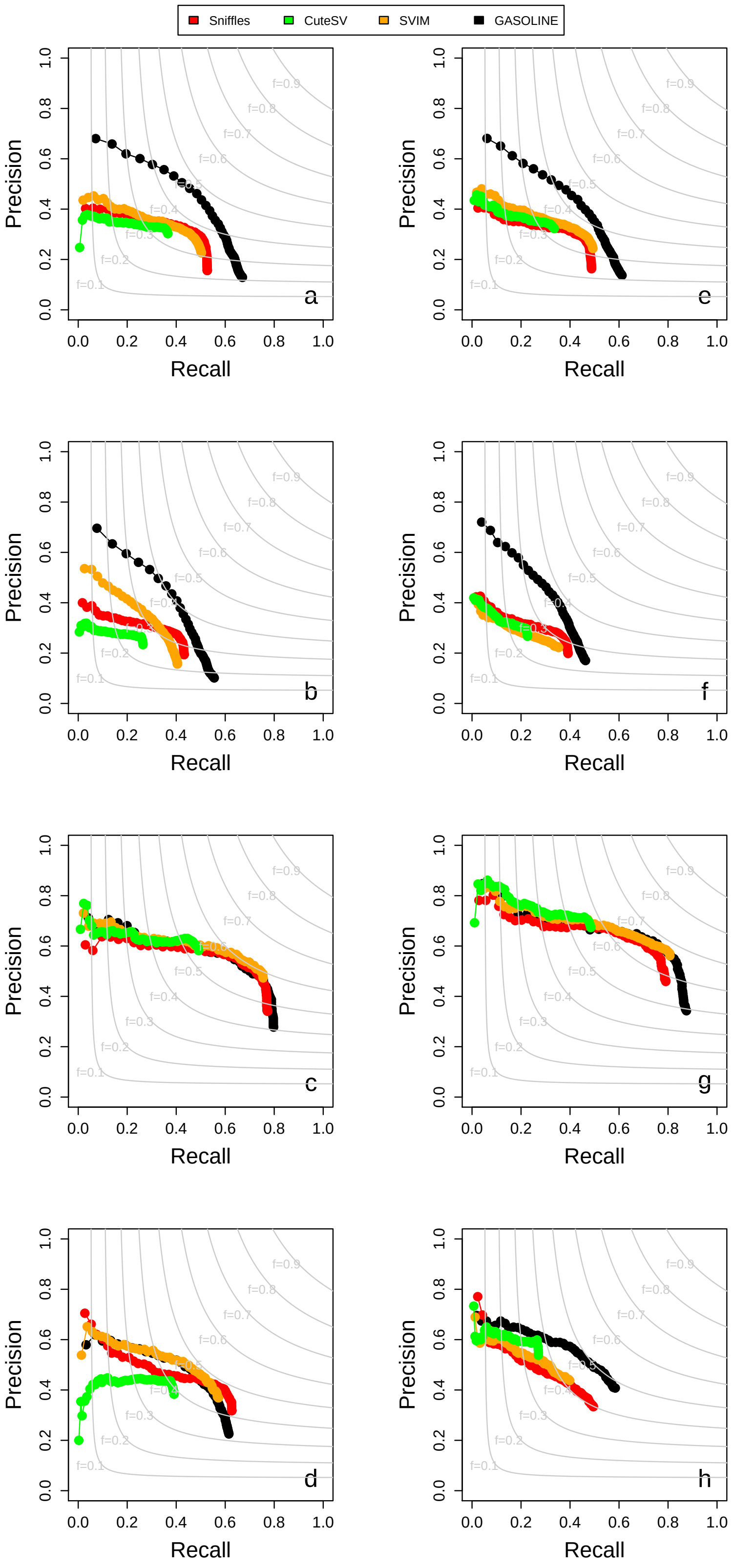


Figure 8: Precision-Recall curves of GASOLINE and the other three tools as a function of number of supporting reads for NA24385 ONT data downsampled at 20x. Panels (a-d) show the results for minimap2 alignment and (e-h) for NGMLR. Panels (a, e) for small deletions (< 500 bp), (b, f) for small insertions (< 500 bp), (a, e) for large deletions (> 500 bp), (a, e) for large insertions (> 500 bp). The curves in panels were obtained by ordering all the SVs as a function of number of supporting reads and calculating precision and recall including SVs with decreasing number of reads.

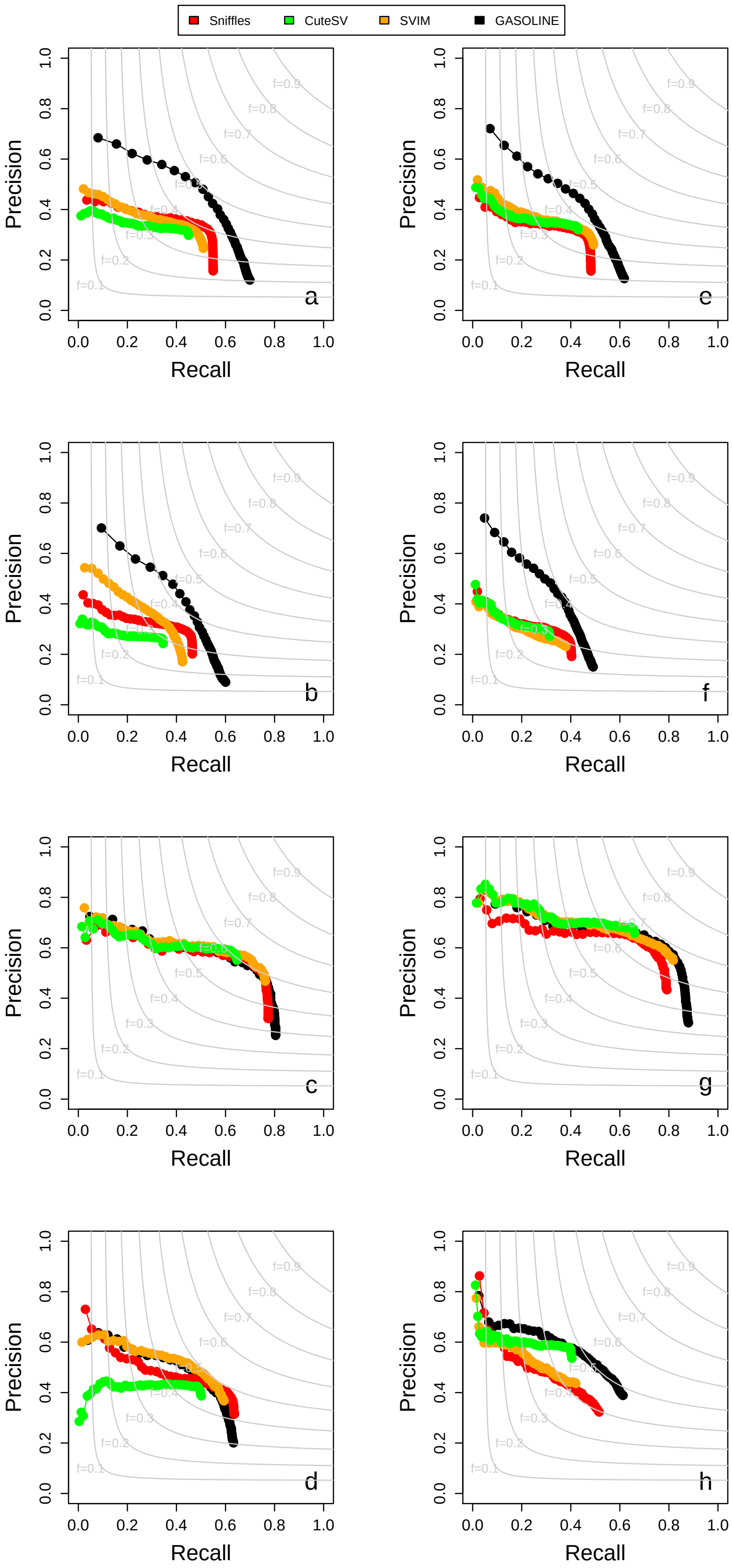


Figure 9: Precision-Recall curves of GASOLINE and the other three tools as a function of number of supporting reads for NA24385 ONT data downsampled at 25x. Panels (a-d) show the results for minimap2 alignment and (e-h) for NGMLR. Panels (a, e) for small deletions (< 500 bp), (b, f) for small insertions (< 500 bp), (a, e) for large deletions (> 500 bp), (a, e) for large insertions (> 500 bp). The curves in panels were obtained by ordering all the SVs as a function of number of supporting reads and calculating precision and recall including SVs with decreasing number of reads.

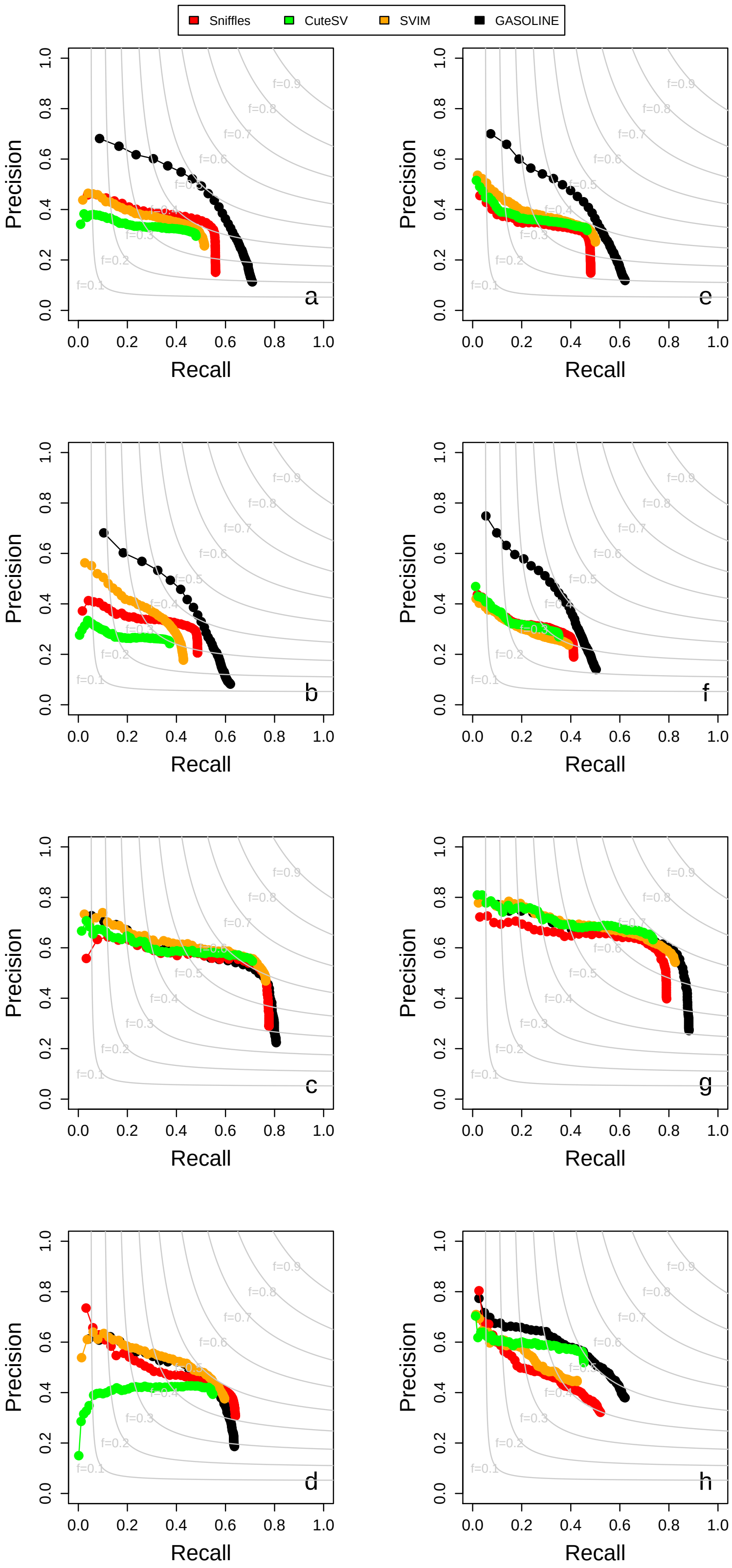


Figure 10: Precision-Recall curves of GASOLINE and the other three tools as a function of number of supporting reads for NA24385 ONT data downsampled at 30x. Panels (a-d) show the results for minimap2 alignment and (e-h) for NGMLR. Panels (a, e) for small deletions (< 500 bp), (b, f) for small insertions (< 500 bp), (a, e) for large deletions (> 500 bp), (a, e) for large insertions (> 500 bp). The curves in panels were obtained by ordering all the SVs as a function of number of supporting reads and calculating precision and recall including SVs with decreasing number of reads.

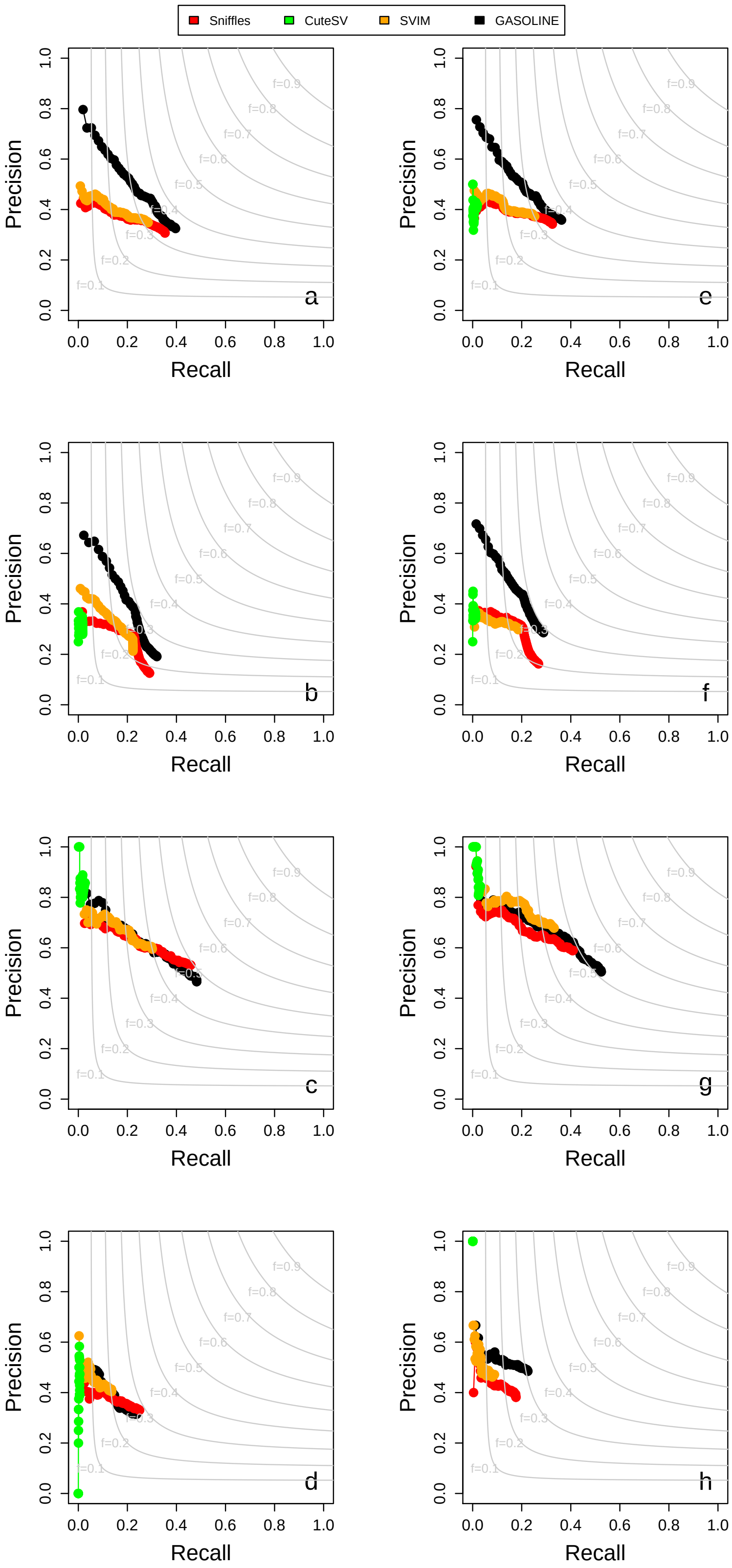


Figure 11: Precision-Recall curves of GASOLINE and the other three tools as a function of number of supporting reads for NA24385 PacBio data downsampled at 5x. Panels (a-d) show the results for minimap2 alignment and (e-h) for NGMLR. Panels (a, e) for small deletions (< 500 bp), (b, f) for small insertions (< 500 bp), (a, e) for large deletions (> 500 bp), (a, e) for large insertions (> 500 bp). The curves in panels were obtained by ordering all the SVs as a function of number of supporting reads and calculating precision and recall including SVs with decreasing number of reads.

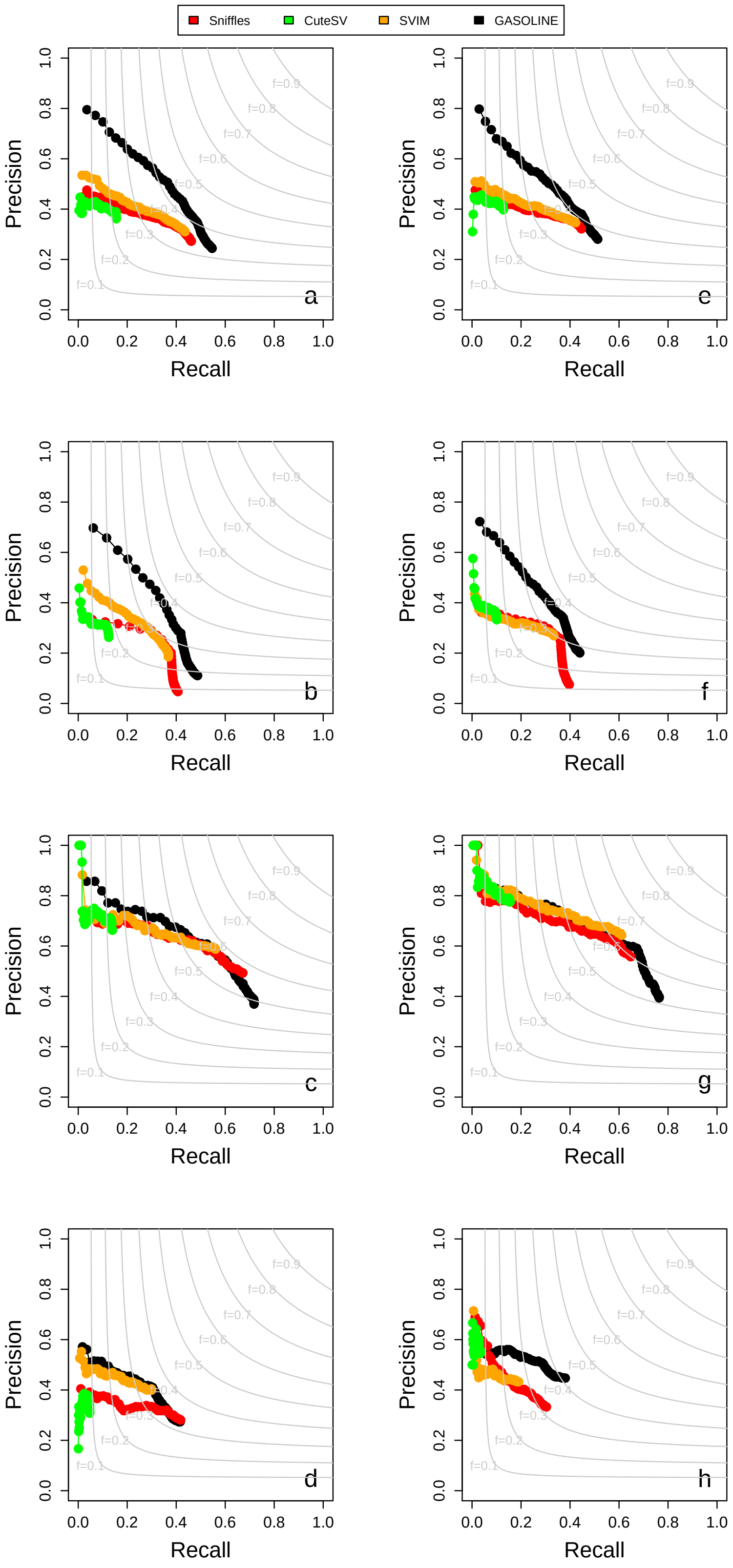


Figure 12: Precision-Recall curves of GASOLINE and the other three tools as a function of number of supporting reads for NA24385 PacBio data downsampled at 10x. Panels (a-d) show the results for minimap2 alignment and (e-h) for NGMLR. Panels (a, e) for small deletions (< 500 bp), (b, f) for small insertions (< 500 bp), (a, e) for large deletions (> 500 bp), (a, e) for large insertions (> 500 bp). The curves in panels were obtained by ordering all the SVs as a function of number of supporting reads and calculating precision and recall including SVs with decreasing number of reads.

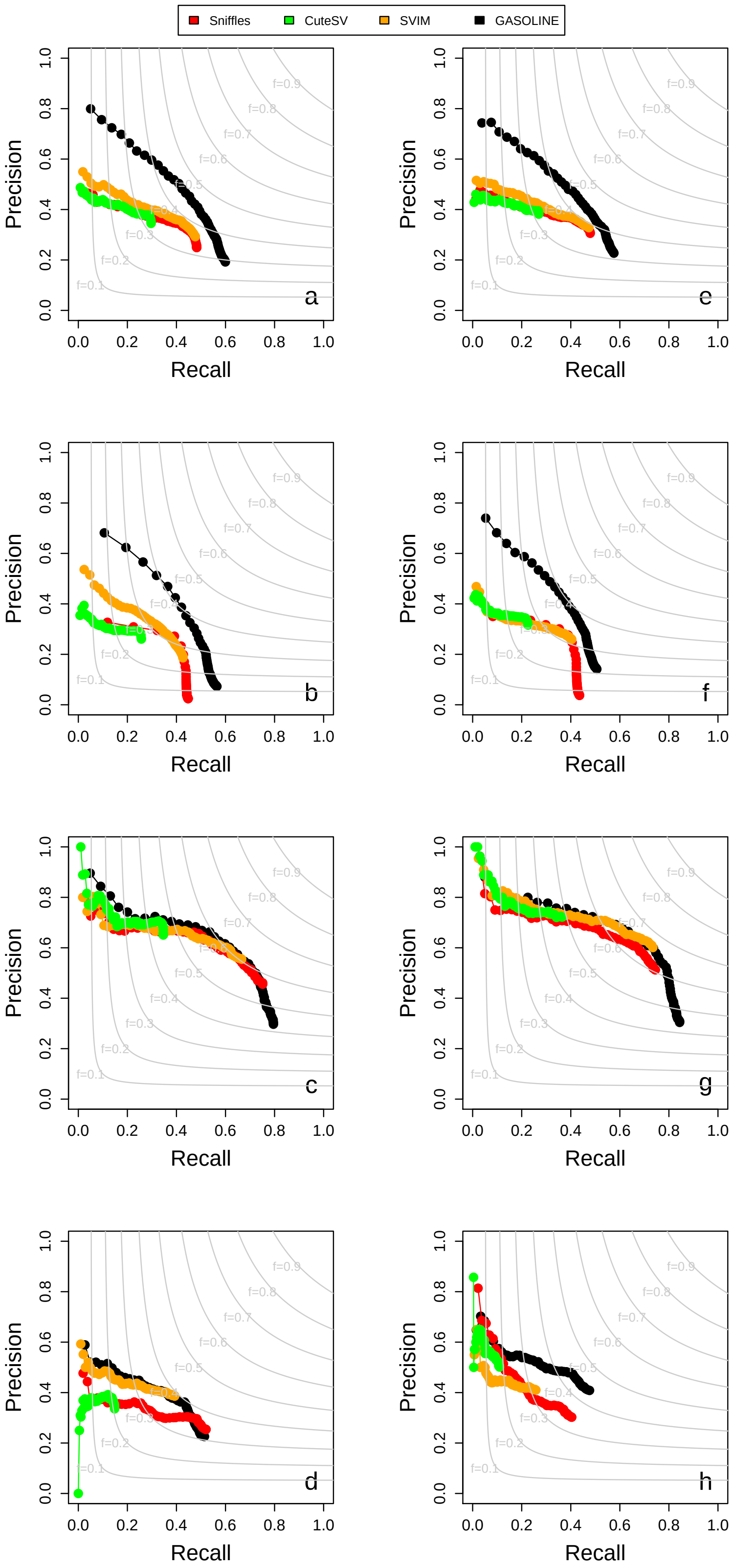


Figure 13: Precision-Recall curves of GASOLINE and the other three tools as a function of number of supporting reads for NA24385 PacBio data downsampled at 15x. Panels (a-d) show the results for minimap2 alignment and (e-h) for NGMLR. Panels (a, e) for small deletions (< 500 bp), (b, f) for small insertions (< 500 bp), (a, e) for large deletions (> 500 bp), (a, e) for large insertions (> 500 bp). The curves in panels were obtained by ordering all the SVs as a function of number of supporting reads and calculating precision and recall including SVs with decreasing number of reads.

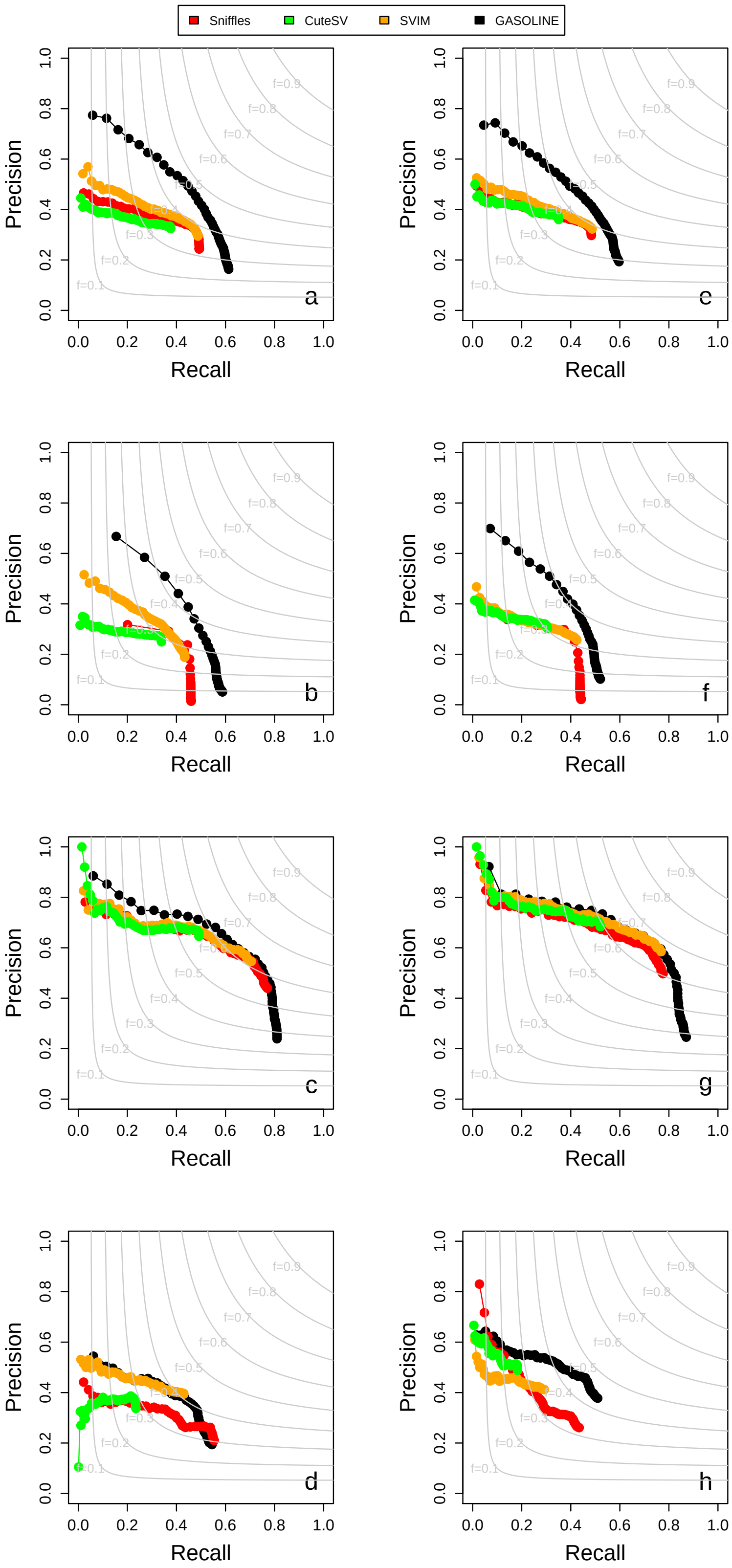


Figure 14: Precision-Recall curves of GASOLINE and the other three tools as a function of number of supporting reads for NA24385 PacBio data downsampled at 20x. Panels (a-d) show the results for minimap2 alignment and (e-h) for NGMLR. Panels (a, e) for small deletions (< 500 bp), (b, f) for small insertions (< 500 bp), (a, e) for large deletions (> 500 bp), (a, e) for large insertions (> 500 bp). The curves in panels were obtained by ordering all the SVs as a function of number of supporting reads and calculating precision and recall including SVs with decreasing number of reads.

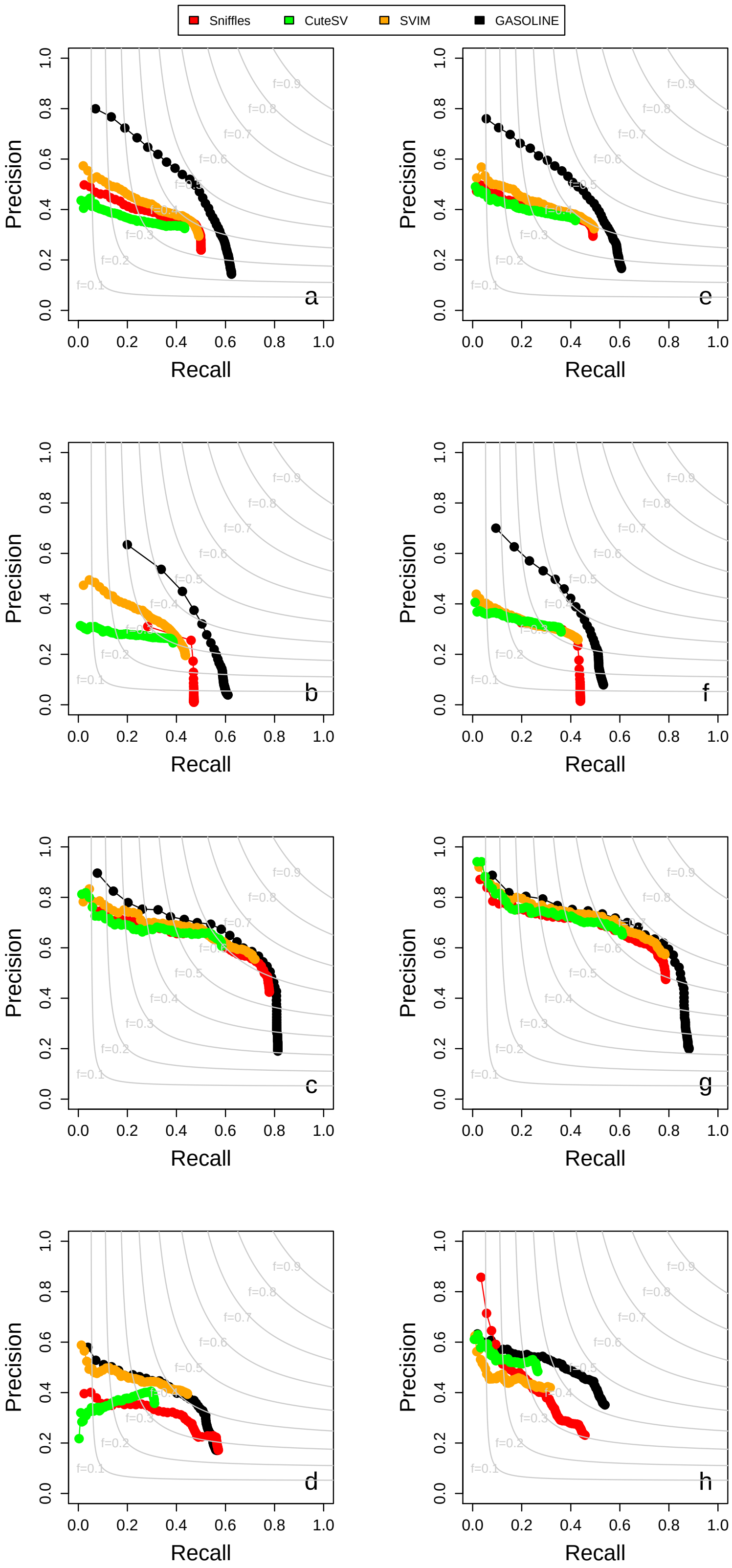


Figure 15: Precision-Recall curves of GASOLINE and the other three tools as a function of number of supporting reads for NA24385 PacBio data downsampled at 25x. Panels (a-d) show the results for minimap2 alignment and (e-h) for NGMLR. Panels (a, e) for small deletions (< 500 bp), (b, f) for small insertions (< 500 bp), (a, e) for large deletions (> 500 bp), (a, e) for large insertions (> 500 bp). The curves in panels were obtained by ordering all the SVs as a function of number of supporting reads and calculating precision and recall including SVs with decreasing number of reads.

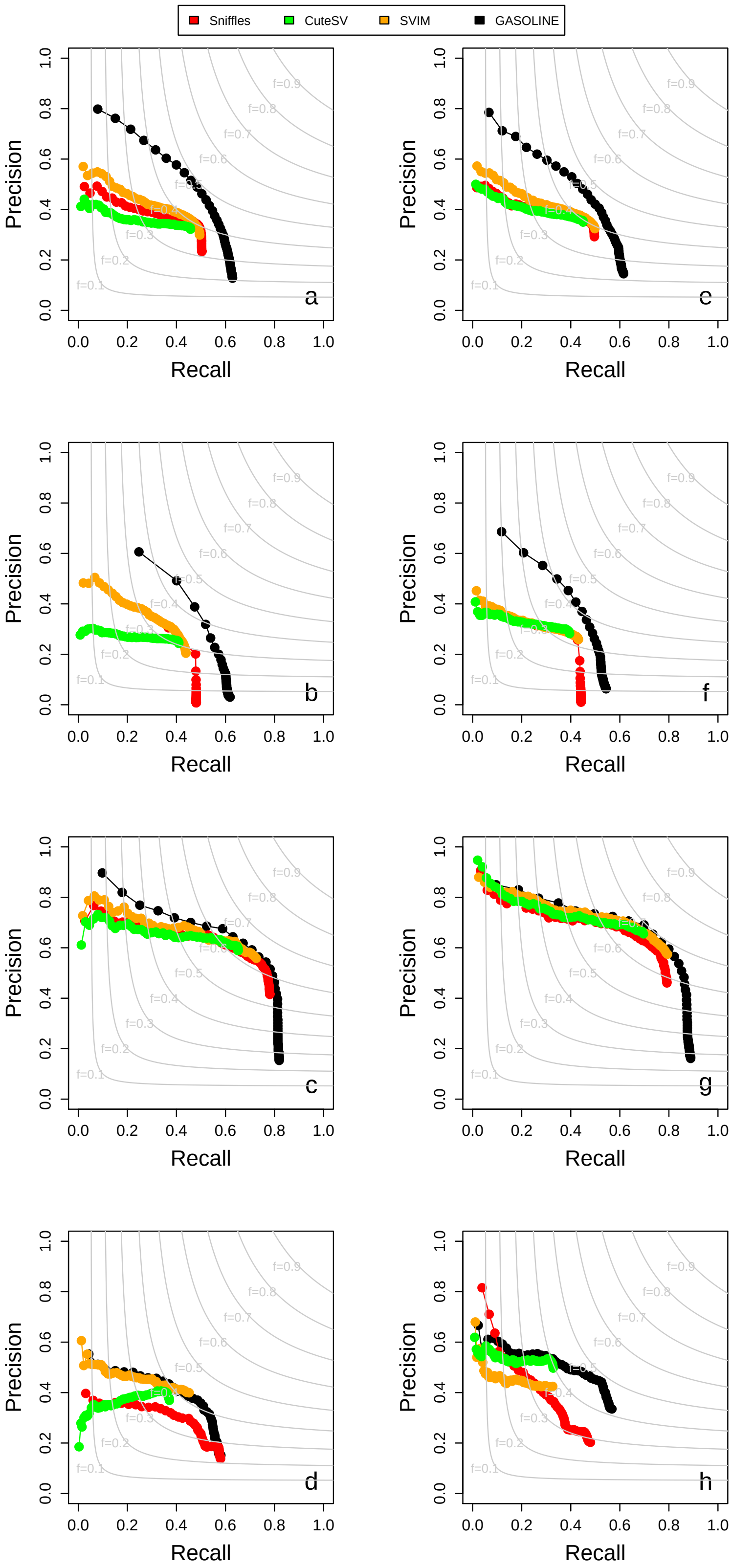


Figure 16: Precision-Recall curves of GASOLINE and the other three tools as a function of number of supporting reads for NA24385 PacBio data downsampled at 30x. Panels (a-d) show the results for minimap2 alignment and (e-h) for NGMLR. Panels (a, e) for small deletions (< 500 bp), (b, f) for small insertions (< 500 bp), (a, e) for large deletions (> 500 bp), (a, e) for large insertions (> 500 bp). The curves in panels were obtained by ordering all the SVs as a function of number of supporting reads and calculating precision and recall including SVs with decreasing number of reads.

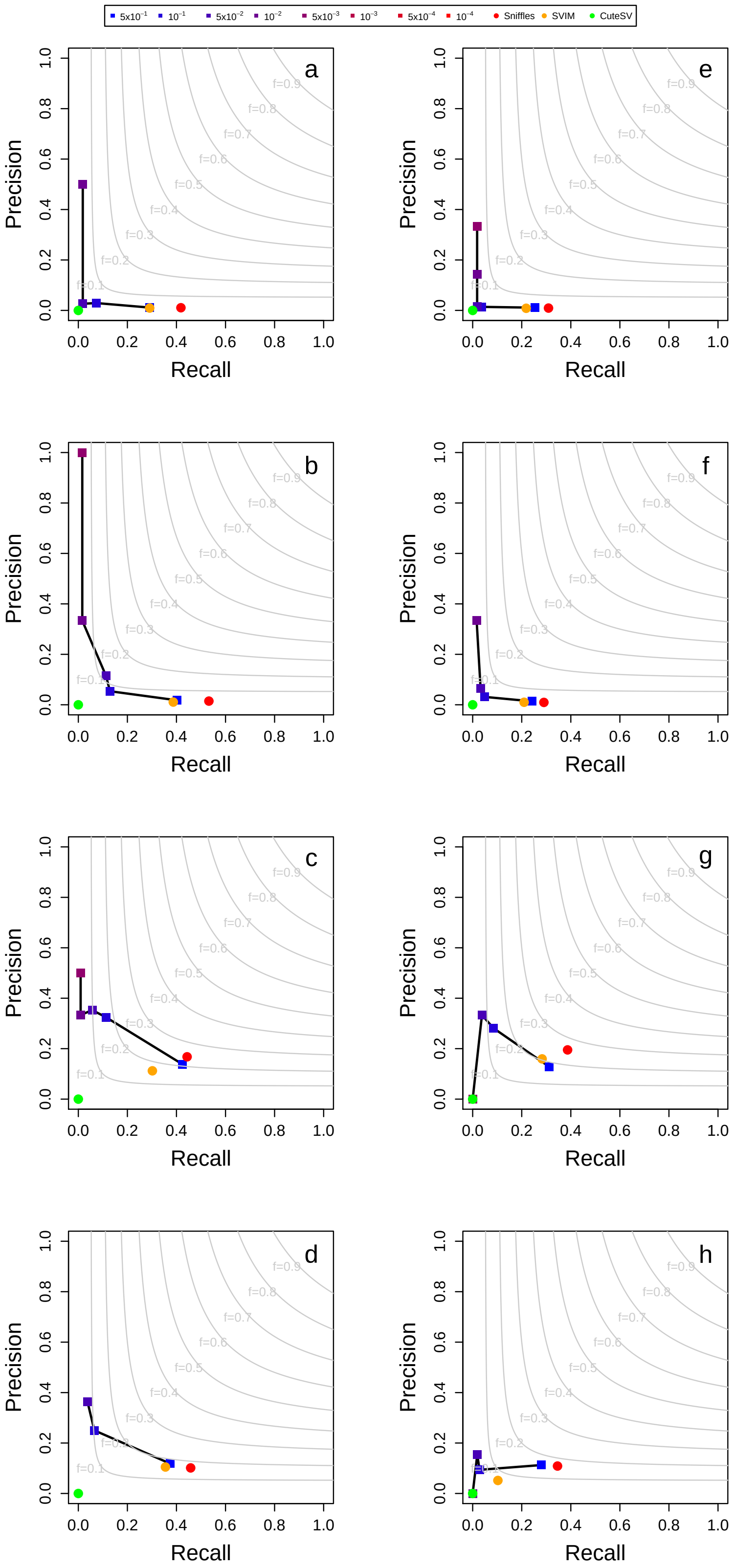


Figure 17: Precision and recall obtained by GASOLINE and the other three tools in the detection simulated somatic SVs from NA24385 ONT data downsampled at 5x. Panels (a-d) show the results for minimap2 alignment and (e-h) for NGMLR. Panels (a, e) for small deletions (< 500 bp), (b, f) for small insertions (< 500 bp), (a, e) for large deletions (> 500 bp), (a, e) for large insertions (> 500 bp). The results for GASOLINE were reported for different somatic p-value thresholds ($5x10^{-1}$, $1x10^{-1}$, $5x10^{-2}$, $1x10^{-2}$, $5x10^{-3}$, $1x10^{-3}$, $5x10^{-4}$, $1x10^{-4}$).

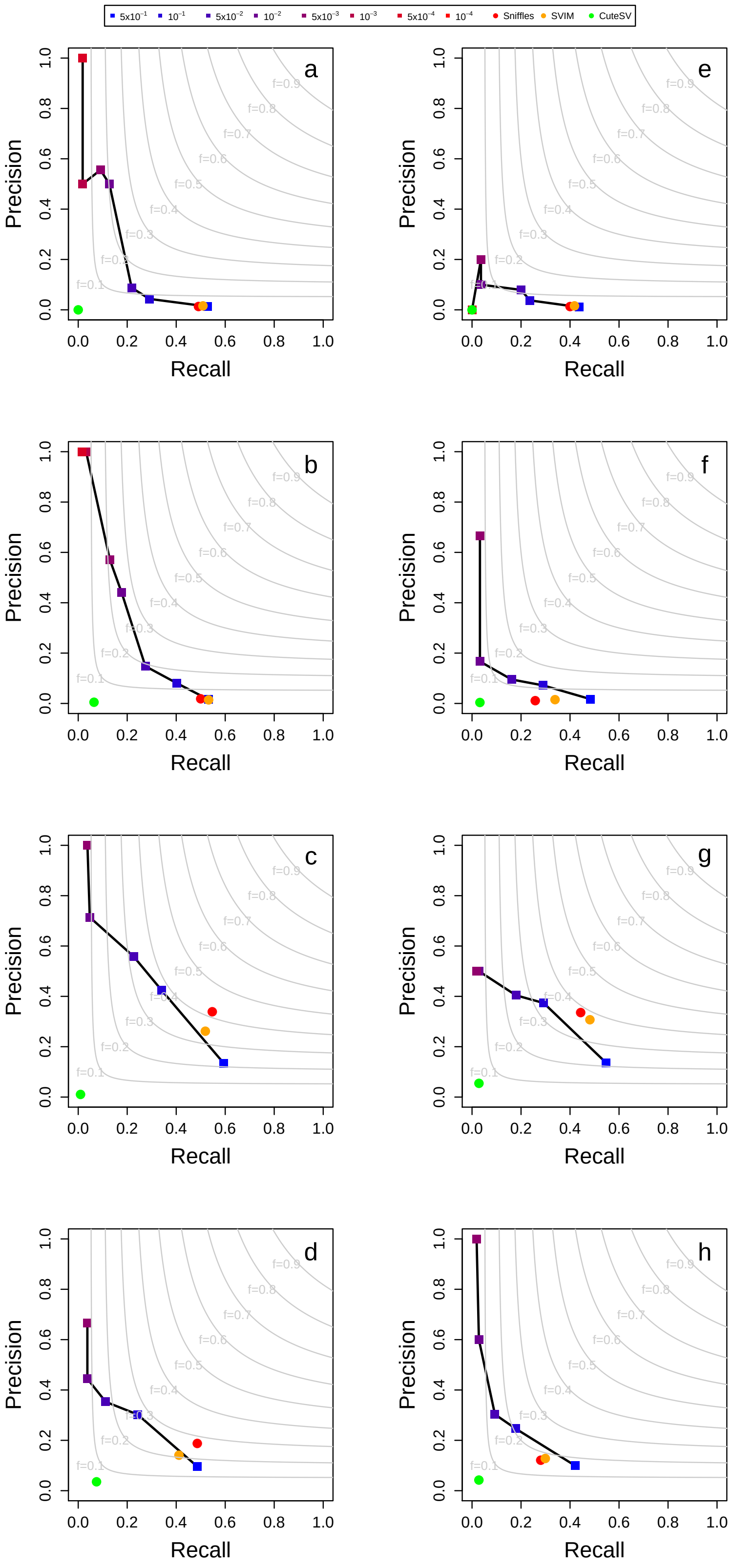


Figure 18: Precision and recall obtained by GASOLINE and the other three tools in the detection simulated somatic SVs from NA24385 ONT data downsampled at 10x. Panels (a-d) show the results for minimap2 alignment and (e-h) for NGMLR. Panels (a, e) for small deletions (< 500 bp), (b, f) for small insertions (< 500 bp), (a, e) for large deletions (> 500 bp), (a, e) for large insertions (> 500 bp). The results for GASOLINE were reported for different somatic p-value thresholds ($5x10^{-1}$, $1x10^{-1}$, $5x10^{-2}$, $1x10^{-2}$, $5x10^{-3}$, $1x10^{-3}$, $5x10^{-4}$, $1x10^{-4}$).

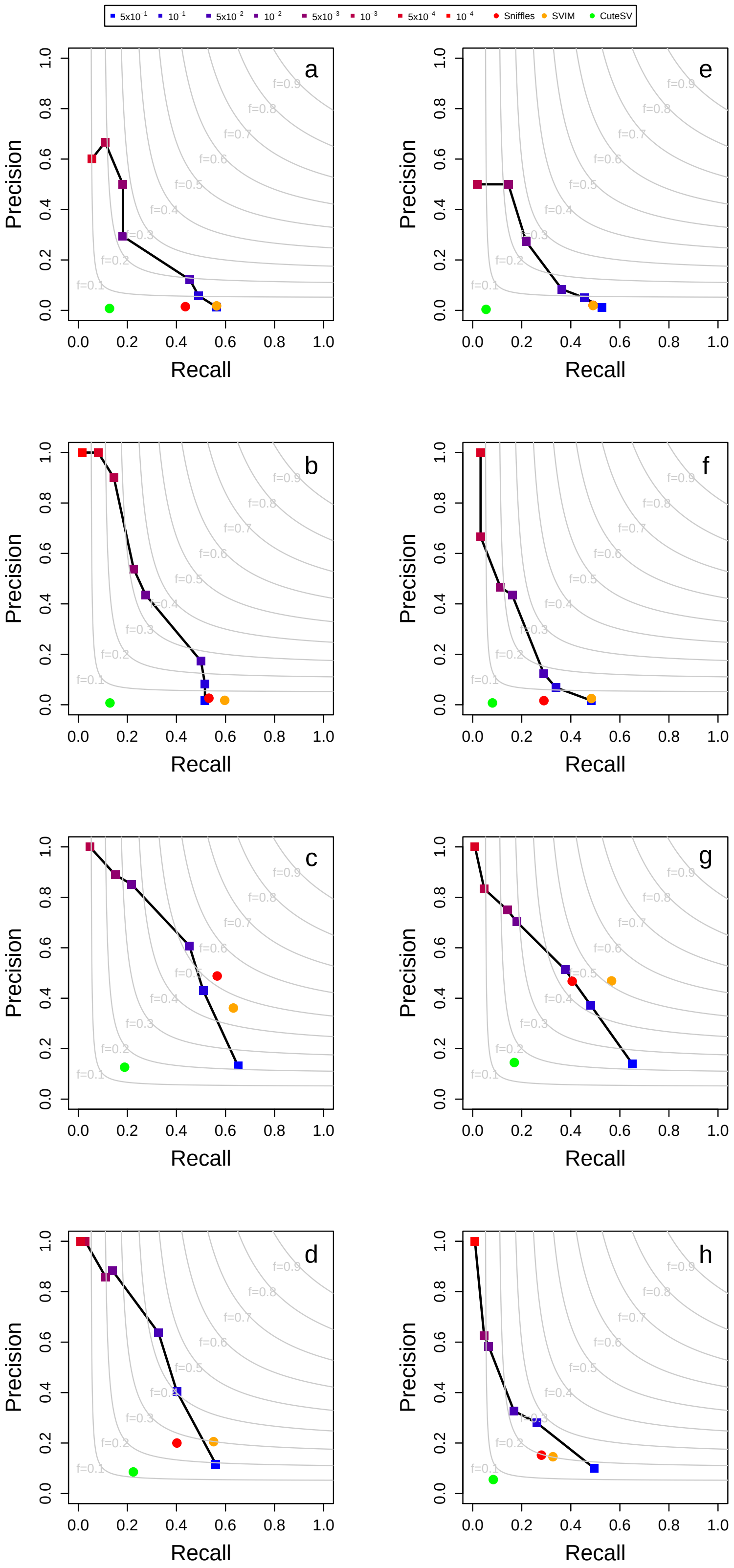


Figure 19: Precision and recall obtained by GASOLINE and the other three tools in the detection simulated somatic SVs from NA24385 ONT data downsampled at 15x. Panels (a-d) show the results for minimap2 alignment and (e-h) for NGMLR. Panels (a, e) for small deletions (< 500 bp), (b, f) for small insertions (< 500 bp), (a, e) for large deletions (> 500 bp), (a, e) for large insertions (> 500 bp). The results for GASOLINE were reported for different somatic p-value thresholds ($5x10^{-1}$, $1x10^{-1}$, $5x10^{-2}$, $1x10^{-2}$, $5x10^{-3}$, $1x10^{-3}$, $5x10^{-4}$, $1x10^{-4}$).

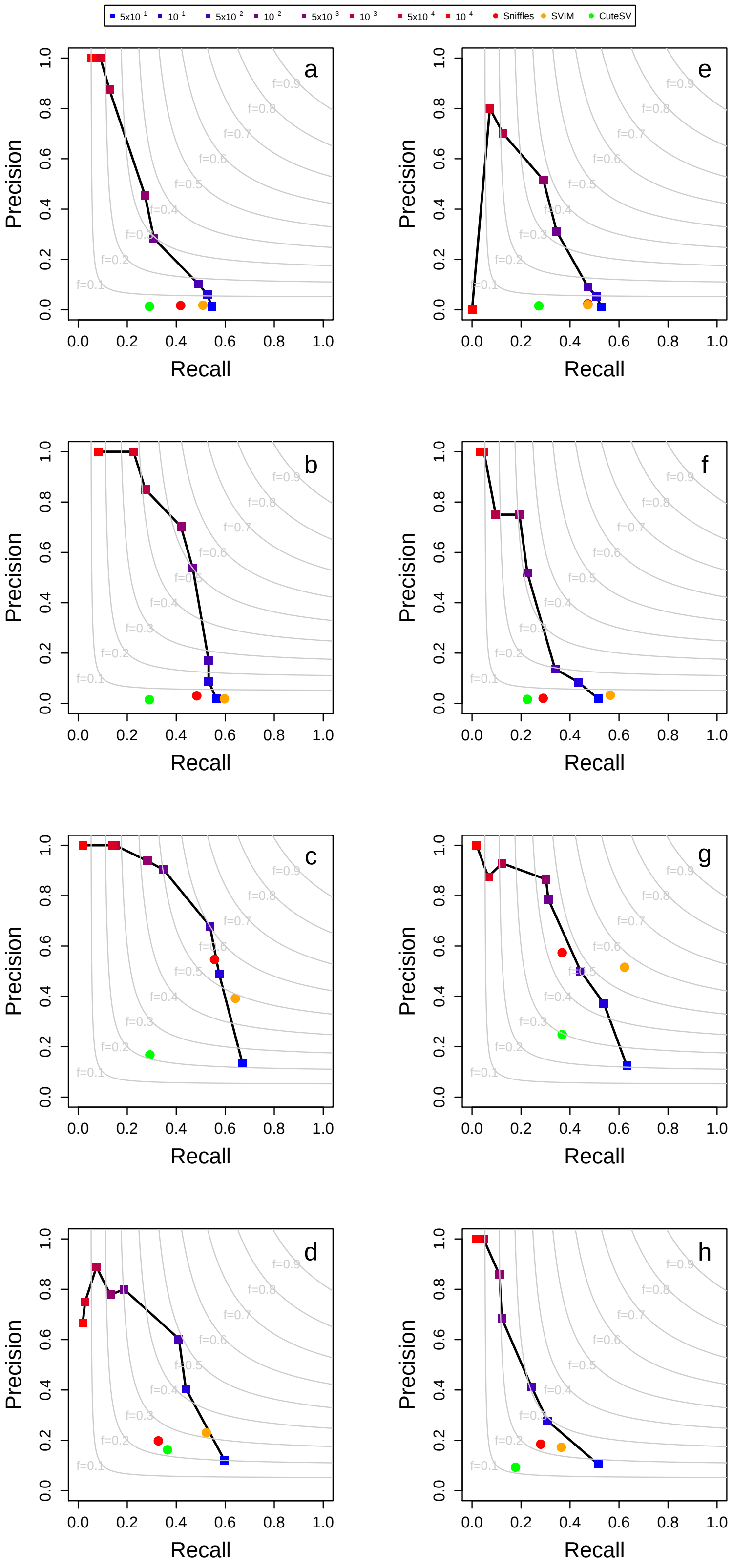


Figure 20: Precision and recall obtained by GASOLINE and the other three tools in the detection simulated somatic SVs from NA24385 ONT data downsampled at 20x. Panels (a-d) show the results for minimap2 alignment and (e-h) for NGMLR. Panels (a, e) for small deletions (< 500 bp), (b, f) for small insertions (< 500 bp), (a, e) for large deletions (> 500 bp), (a, e) for large insertions (> 500 bp). The results for GASOLINE were reported for different somatic p-value thresholds ($5x10^{-1}$, $1x10^{-1}$, $5x10^{-2}$, $1x10^{-2}$, $5x10^{-3}$, $1x10^{-3}$, $5x10^{-4}$, $1x10^{-4}$).

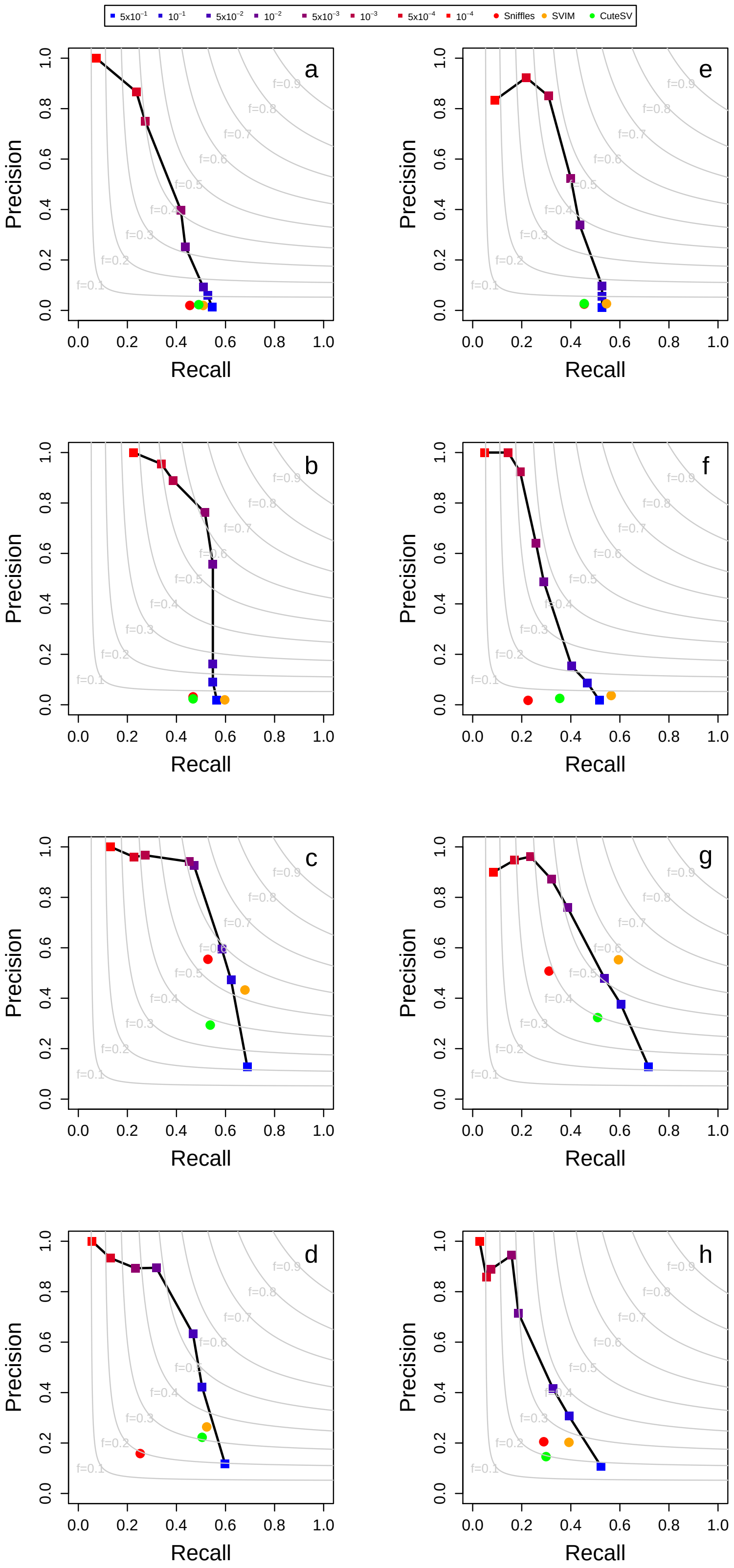


Figure 21: Precision and recall obtained by GASOLINE and the other three tools in the detection simulated somatic SVs from NA24385 ONT data downsampled at 25x. Panels (a-d) show the results for minimap2 alignment and (e-h) for NGMLR. Panels (a, e) for small deletions ($<$ 500 bp), (b, f) for small insertions ($<$ 500 bp), (a, e) for large deletions ($>$ 500 bp), (a, e) for large insertions ($>$ 500 bp). The results for GASOLINE were reported for different somatic p-value thresholds ($5x10^{-1}$, $1x10^{-1}$, $5x10^{-2}$, $1x10^{-2}$, $5x10^{-3}$, $1x10^{-3}$, $5x10^{-4}$, $1x10^{-4}$).

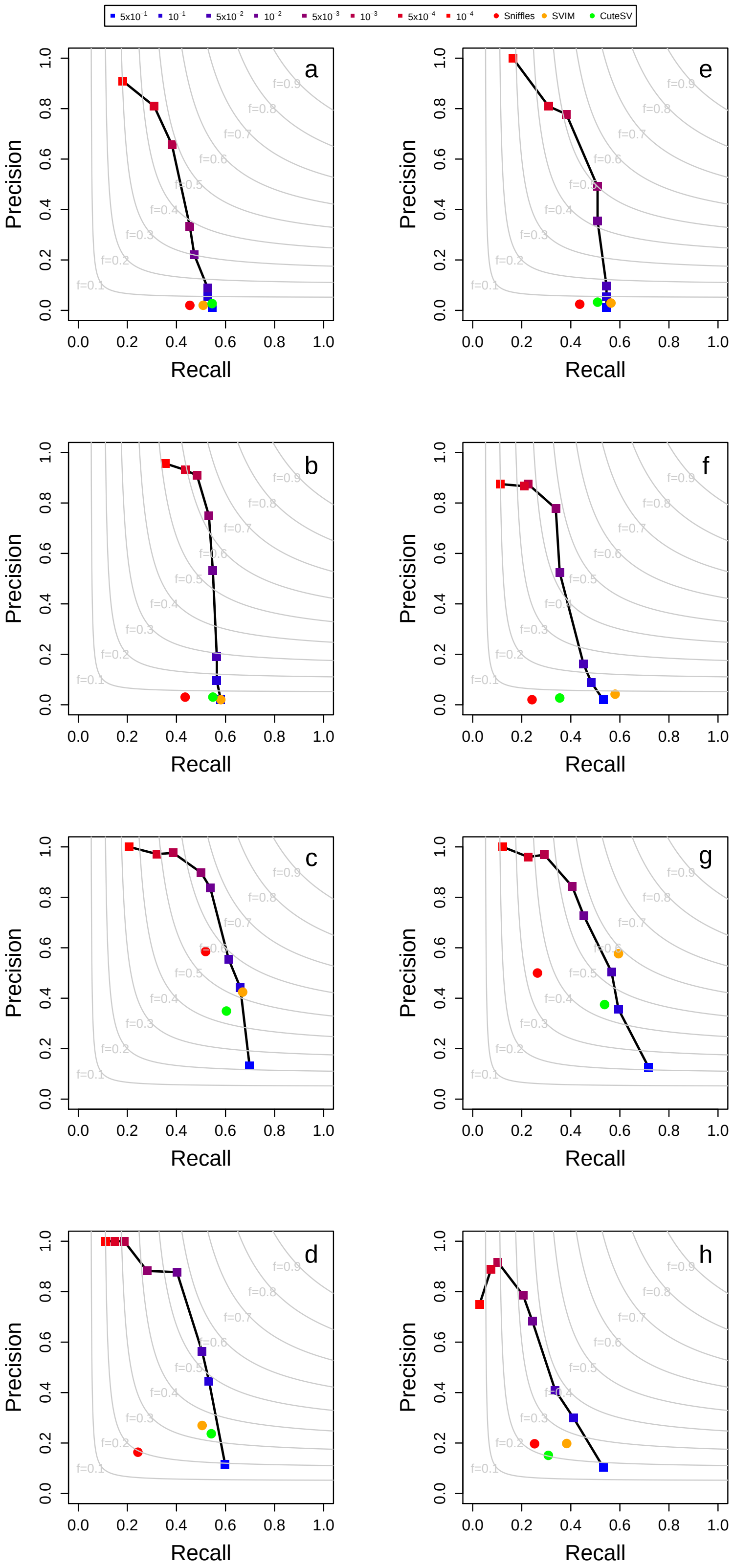


Figure 22: Precision and recall obtained by GASOLINE and the other three tools in the detection simulated somatic SVs from NA24385 ONT data downsampled at 30x. Panels (a-d) show the results for minimap2 alignment and (e-h) for NGMLR. Panels (a, e) for small deletions (< 500 bp), (b, f) for small insertions (< 500 bp), (a, e) for large deletions (> 500 bp), (a, e) for large insertions (> 500 bp). The results for GASOLINE were reported for different somatic p-value thresholds ($5x10^{-1}$, $1x10^{-1}$, $5x10^{-2}$, $1x10^{-2}$, $5x10^{-3}$, $1x10^{-3}$, $5x10^{-4}$, $1x10^{-4}$).

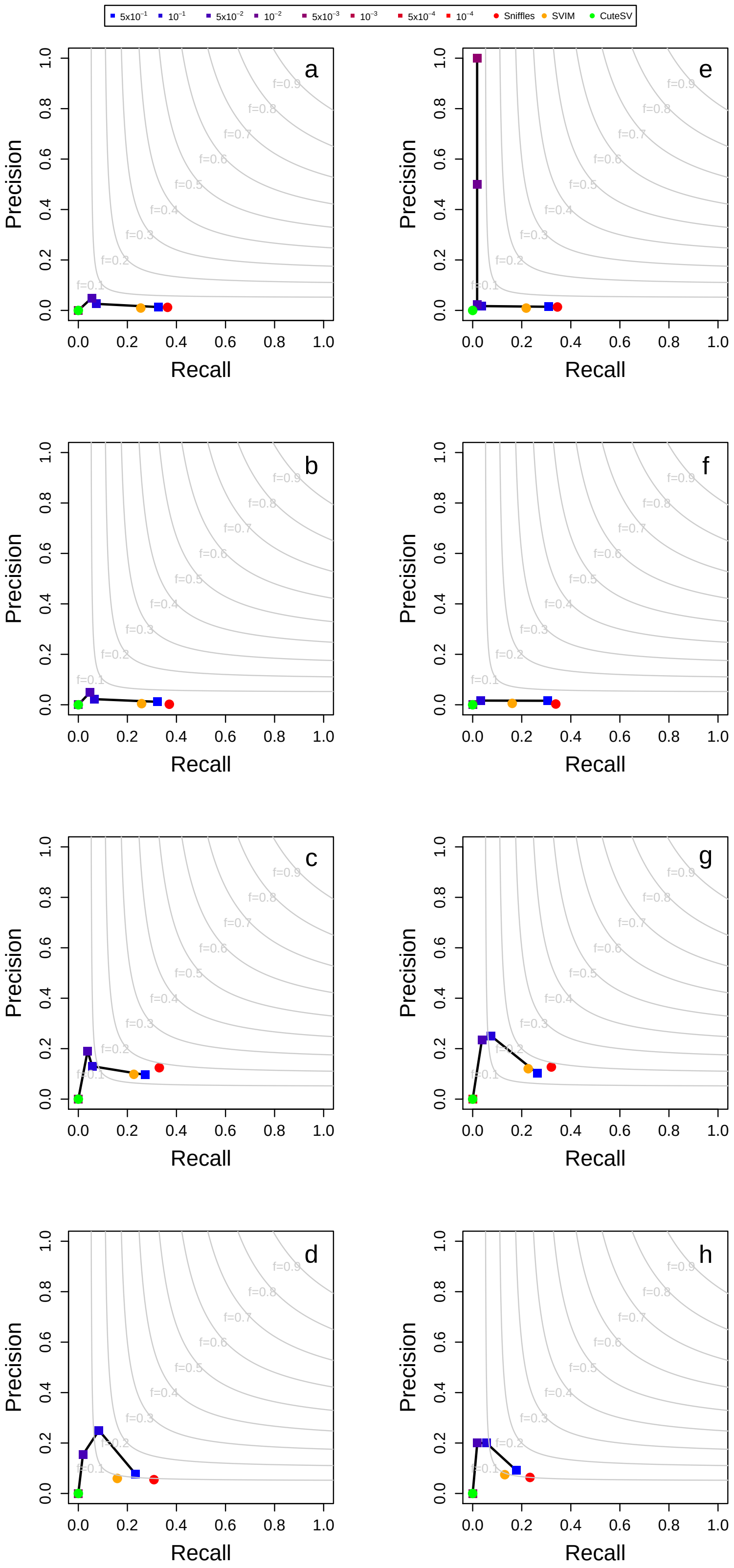


Figure 23: Precision and recall obtained by GASOLINE and the other three tools in the detection simulated somatic SVs from NA24385 PacBio data downsampled at 5x. Panels (a-d) show the results for minimap2 alignment and (e-h) for NGMLR. Panels (a, e) for small deletions (< 500 bp), (b, f) for small insertions (< 500 bp), (a, e) for large deletions (> 500 bp), (a, e) for large insertions (> 500 bp). The results for GASOLINE were reported for different somatic p-value thresholds ($5x10^{-1}$, $1x10^{-1}$, $5x10^{-2}$, $1x10^{-2}$, $5x10^{-3}$, $1x10^{-3}$, $5x10^{-4}$, $1x10^{-4}$).

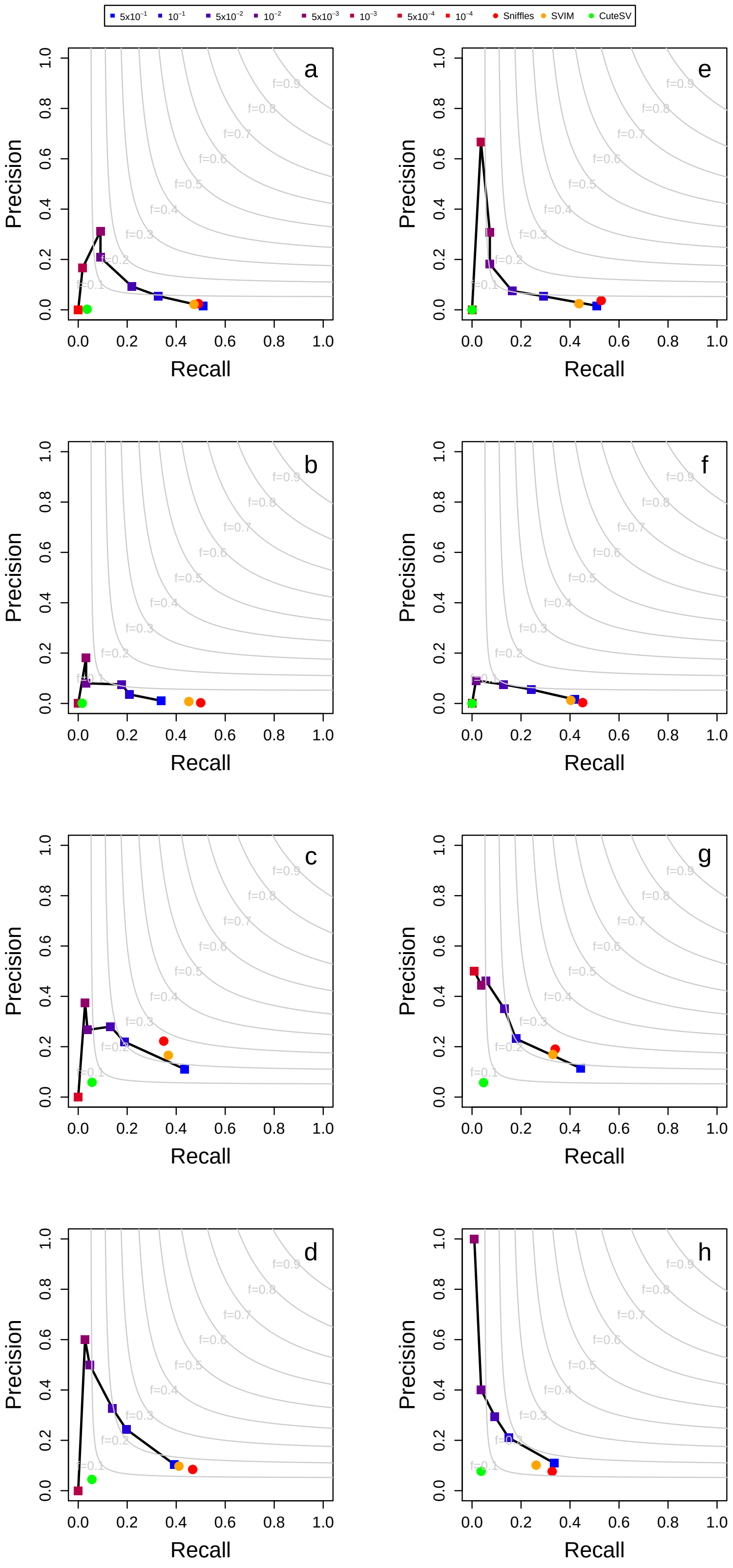


Figure 24: Precision and recall obtained by GASOLINE and the other three tools in the detection simulated somatic SVs from NA24385 PacBio data downsampled at 10x. Panels (a-d) show the results for minimap2 alignment and (e-h) for NGMLR. Panels (a, e) for small deletions (< 500 bp), (b, f) for small insertions (< 500 bp), (a, e) for large deletions (> 500 bp), (a, e) for large insertions (> 500 bp). The results for GASOLINE were reported for different somatic p-value thresholds ($5x10^{-1}$, $1x10^{-1}$, $5x10^{-2}$, $1x10^{-2}$, $5x10^{-3}$, $1x10^{-3}$, $5x10^{-4}$, $1x10^{-4}$).

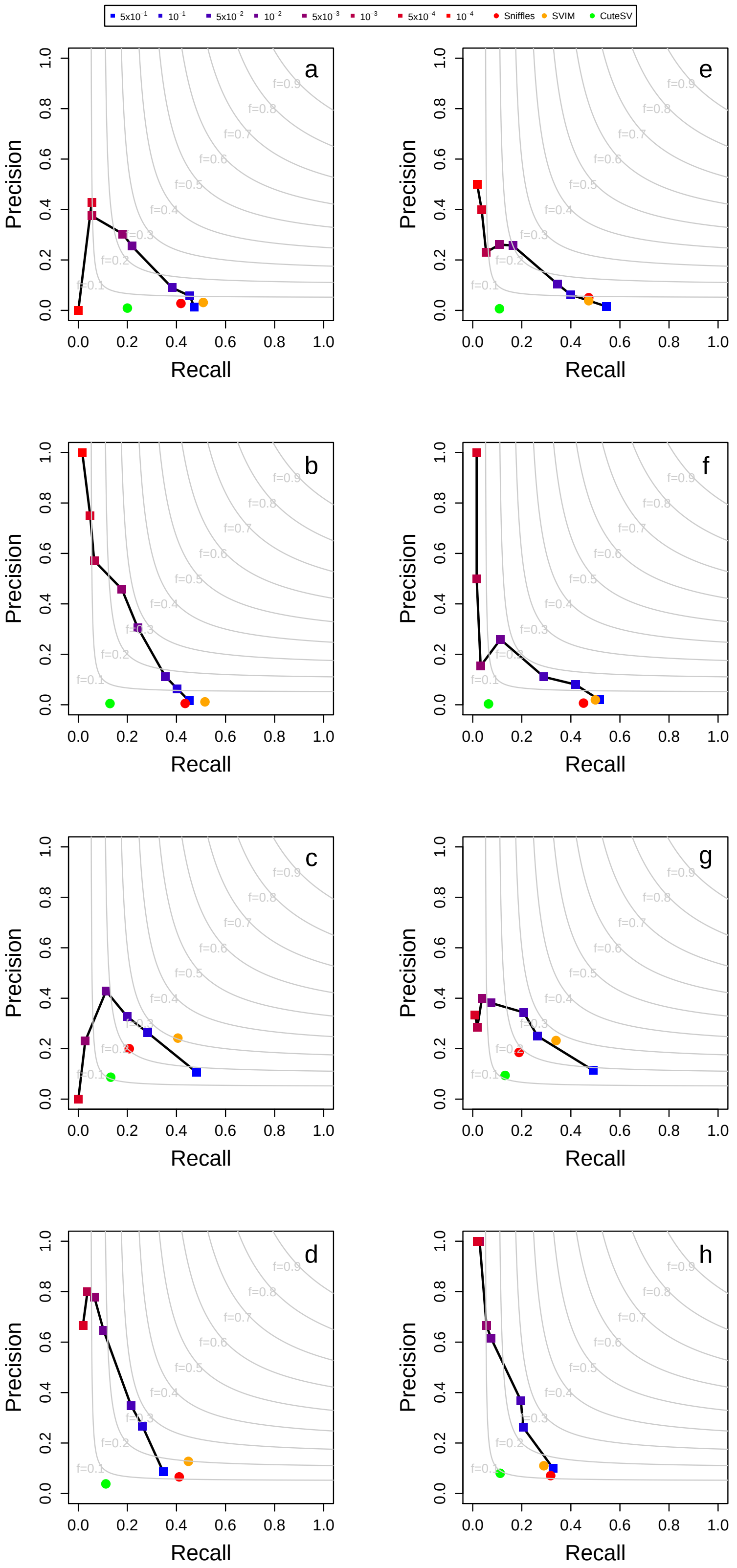


Figure 25: Precision and recall obtained by GASOLINE and the other three tools in the detection simulated somatic SVs from NA24385 PacBio data downsampled at 15x. Panels (a-d) show the results for minimap2 alignment and (e-h) for NGMLR. Panels (a, e) for small deletions (< 500 bp), (b, f) for small insertions (< 500 bp), (a, e) for large deletions (> 500 bp), (a, e) for large insertions (> 500 bp). The results for GASOLINE were reported for different somatic p-value thresholds ($5x10^{-1}$, $1x10^{-1}$, $5x10^{-2}$, $1x10^{-2}$, $5x10^{-3}$, $1x10^{-3}$, $5x10^{-4}$, $1x10^{-4}$).

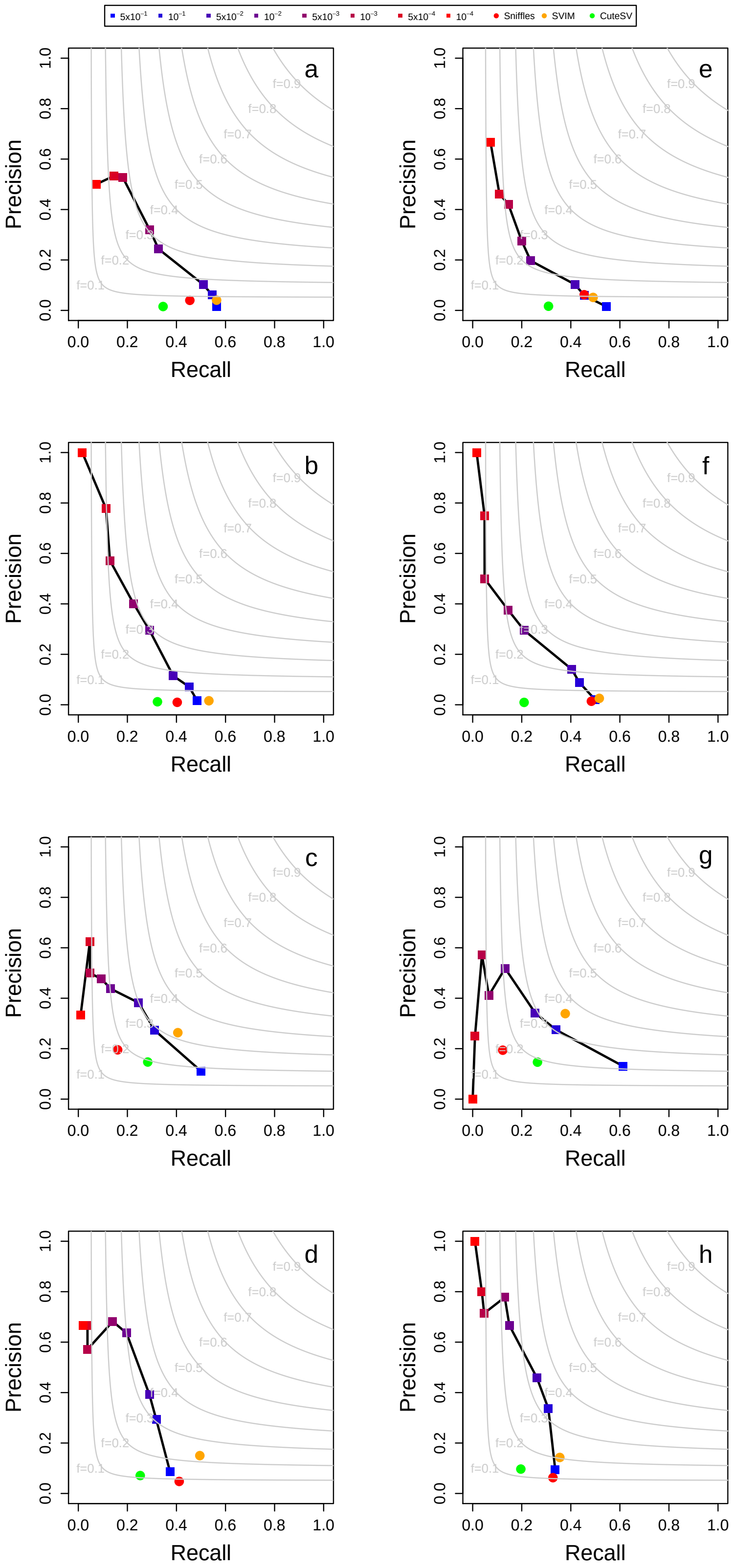


Figure 26: Precision and recall obtained by GASOLINE and the other three tools in the detection simulated somatic SVs from NA24385 PacBio data downsampled at 20x. Panels (a-d) show the results for minimap2 alignment and (e-h) for NGMLR. Panels (a, e) for small deletions (< 500 bp), (b, f) for small insertions (< 500 bp), (a, e) for large deletions (> 500 bp), (a, e) for large insertions (> 500 bp). The results for GASOLINE were reported for different somatic p-value thresholds ($5x10^{-1}$, $1x10^{-1}$, $5x10^{-2}$, $1x10^{-2}$, $5x10^{-3}$, $1x10^{-3}$, $5x10^{-4}$, $1x10^{-4}$).

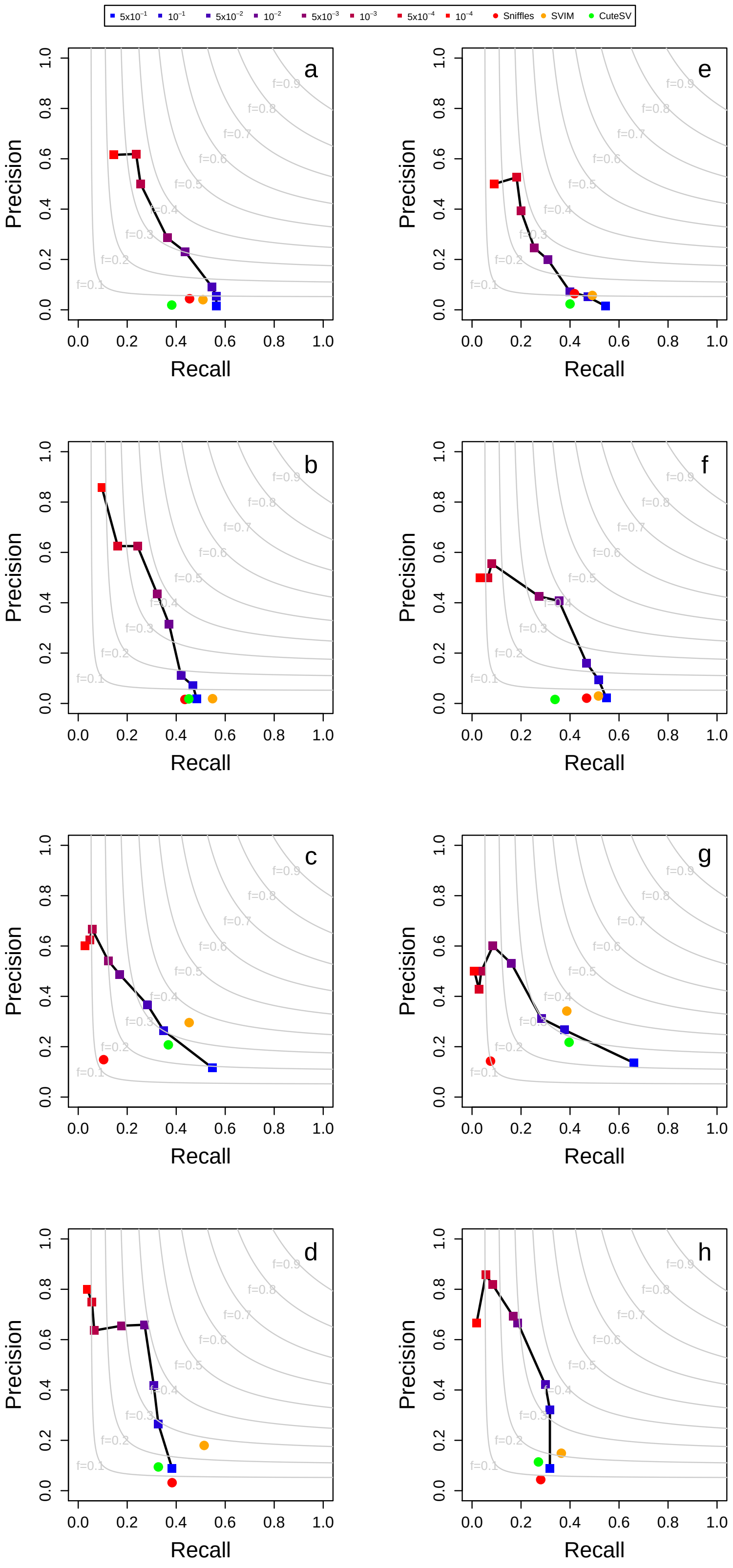


Figure 27: Precision and recall obtained by GASOLINE and the other three tools in the detection simulated somatic SVs from NA24385 PacBio data downsampled at 25x. Panels (a-d) show the results for minimap2 alignment and (e-h) for NGMLR. Panels (a, e) for small deletions (< 500 bp), (b, f) for small insertions (< 500 bp), (a, e) for large deletions (> 500 bp), (a, e) for large insertions (> 500 bp). The results for GASOLINE were reported for different somatic p-value thresholds (5x$10^{-1}$, 1x$10^{-1}$, 5x$10^{-2}$, 1x$10^{-2}$, 5x$10^{-3}$, 1x$10^{-3}$, 5x$10^{-4}$, 1x$10^{-4}$).

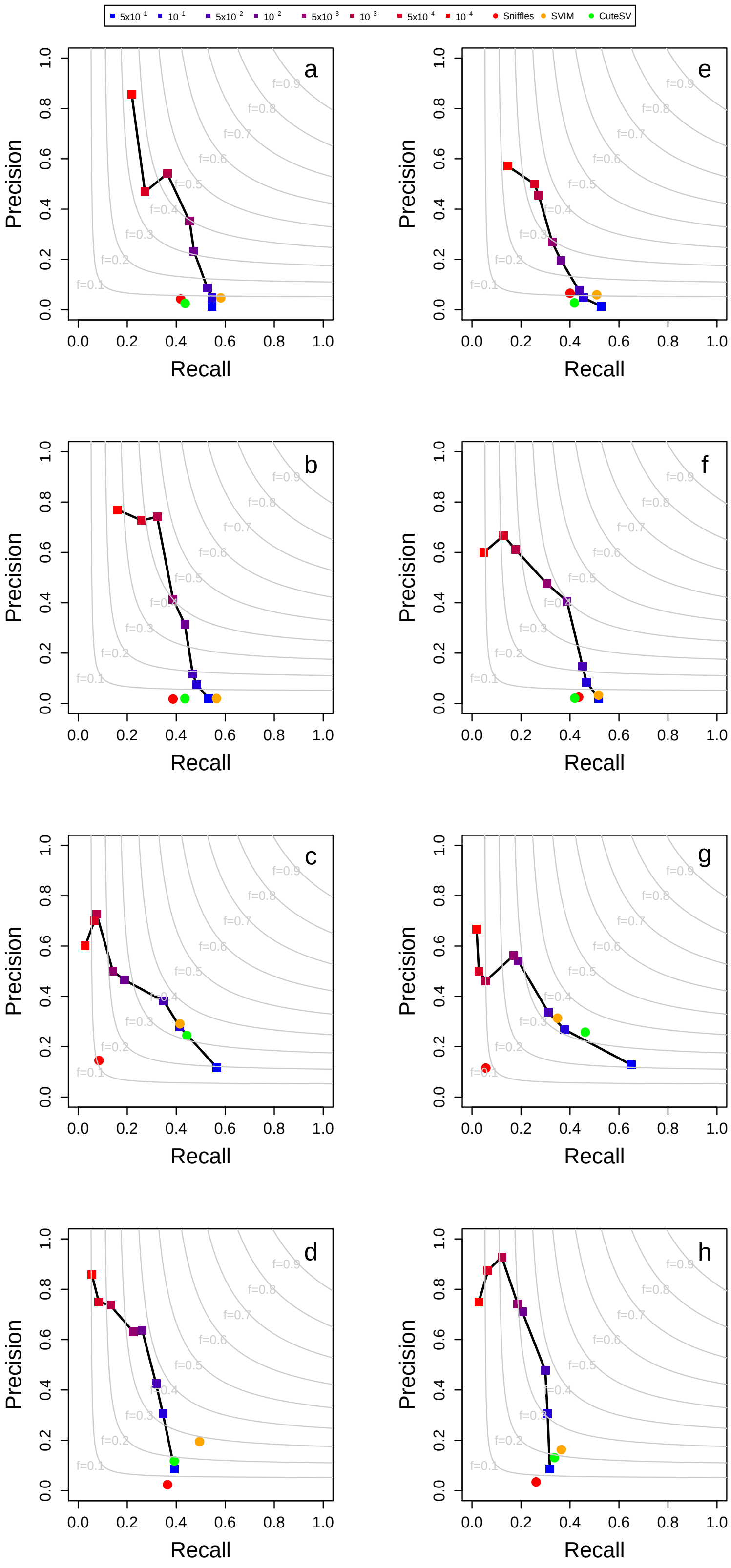


Figure 28: Precision and recall obtained by GASOLINE and the other three tools in the detection simulated somatic SVs from NA24385 PacBio data downsampled at 30x. Panels (a-d) show the results for minimap2 alignment and (e-h) for NGMLR. Panels (a, e) for small deletions (< 500 bp), (b, f) for small insertions (< 500 bp), (a, e) for large deletions (> 500 bp), (a, e) for large insertions (> 500 bp). The results for GASOLINE were reported for different somatic p-value thresholds ($5x10^{-1}$, $1x10^{-1}$, $5x10^{-2}$, $1x10^{-2}$, $5x10^{-3}$, $1x10^{-3}$, $5x10^{-4}$, $1x10^{-4}$).
